# Dis3L2 regulates cellular proliferation through a PI3-Kinase dependent signalling pathway

**DOI:** 10.1101/806109

**Authors:** Benjamin P. Towler, Amy L. Pashler, Hope J. Haime, Katarzyna M. Przybyl, Sandra C. Viegas, Rute G. Matos, Simon J. Morley, Cecilia M. Arraiano, Sarah F. Newbury

**Affiliations:** Medical Research Building, Brighton and Sussex Medical School, University of Sussex, Falmer, Brighton BN1 9PS, UK; Instituto de Tecnologia Química e Biológica António Xavier, Universidade Nova de Lisboa, Av. da República, 2780-157 Oeiras, Portugal; John Maynard Smith Building, School of Life Sciences, University of Sussex, Falmer, Brighton, BN1 9QG, UK

**Keywords:** Tissue growth, *Drosophila*, exoribonucleases, RNA stability, HEK-293T cells, Idgf2

## Abstract

Dis3L2 is a highly conserved 3’-5’ exoribonuclease which is mutated in the human overgrowth disorders Perlman syndrome and Wilms’ tumour. Here, we have generated a new *dis3L*2 null mutant and a UAS-nuclease-dead line in *Drosophila* to demonstrate that the catalytic activity of Dis3L2 is required to control proliferation in imaginal discs. Using RNA-seq on *dis3L2* mutant wing discs, we show that the imaginal disc growth factor Idgf2 is responsible for driving the wing overgrowth observed in *dis3L2* null mutants. IDGFs are conserved proteins homologous to human chitinase-like proteins such as CHI3L1/YKL-40 which are implicated in tissue regeneration as well as cancers including colon cancer and non-small cell lung cancer. We also show that loss of DIS3L2 in human HEK-293T cells results in cell proliferation, illustrating the conservation of this important cell proliferation pathway. Using these human cells we show that loss of DIS3L2 results in an increase in the activity of the PI3-Kinase/AKT signalling pathway, which we subsequently show to also have a role in driving the proliferation phenotype in *Drosophila*. Our work therefore provides the first mechanistic explanation for DIS3L2-induced overgrowth in humans and flies and identifies an ancient proliferation pathway controlled by Dis3L2 to regulate cell proliferation and tissue growth.

## Introduction

The correct size and shape of an organ or tissue within an adult animal depends on a balance between cell division and cell death. The regulation of cell proliferation is also important in regeneration and repair of damaged tissues during wound healing. Co-ordination of tissue growth usually requires release of growth factors to signal between cells of an epithelial layer and communicate between tissues to maintain the correct size and shape of different organs. These signals need to be correctly interpreted by particular RNAs and proteins within cells in each tissue to maintain a balance between proliferation, apoptosis and differentiation. Although the cellular pathways governing uncontrolled cell proliferation as occurs in cancer are well known, the cellular pathways regulating normal, co-ordinated cell proliferation have not been as well studied. At the molecular level, many transcription factors and signalling proteins regulating proliferation (e.g. p53, Wingless/Wnt) have been identified whereas the contribution of RNA degradation to the control of cell proliferation is not well understood.

Dis3L2 is an exoribonuclease from the RNase II/RNB family which degrades RNAs in a 3’-5’ direction. Our previous work, using the model organism *Drosophila*, demonstrated that knockdown of Dis3L2 in the wing imaginal disc results in substantial wing overgrowth due to increased cellular proliferation during the larval stages (1). Remarkably, the proliferation effect seen in *Drosophila* tissue upon Dis3L2 knockdown is similar to that seen in human tissues. Previous work has demonstrated that mutations in *DIS3L2* are associated with Perlman syndrome (2). This syndrome is a congenital overgrowth condition which results in foetal gigantism, abnormal enlargement of organs (e.g. kidneys), facial abnormalities, neurodevelopmental delay and high neonatal mortality. Patients also have a predisposition for Wilms’ tumour (2–4). DIS3L2 is also likely to be directly linked with sporadic Wilms’ tumour as a substantial number of these tumours show partial or complete *DIS3L2* deletion (2,4). In addition, deletions in DIS3L2 have also been associated with skeletal overgrowth (5). Understanding the pathways whereby Dis3L2 regulates cell proliferation in *Drosophila* is therefore likely to shed light not only on the molecular basis of normal cell proliferation but also on these overgrowth diseases.

Dis3L2 is a member of the RNase II/RNB family of 3’-5’ exoribonucleases which is conserved from bacteria through to humans (6). In humans and other higher eukaryotes this family consists of three members: Dis3, Dis3L1 and Dis3L2, whereas *Drosophila melanogaster* encodes two members (Dis3 and Dis3L2) and *Saccharomyces cerevisiae* encodes only one (Dis3). Dis3L2 is cytoplasmic and differs from Dis3 in that it lacks the N-terminal PIN domain (6,7) which confers endoribonucleolytic activity to Dis3 and tethers it to the multicomponent exosome complex (8–10). Dis3L2 also differs from Dis3 in that it shows specificity towards transcripts that are uridylated at their 3’ ends due to the action of terminal uridyltransferases (TUTases) (6,11–15). This specificity is due to structural features of the enzyme in the channel leading to the active site (16).

A number of experiments, using human, mouse, fly and yeast cells have suggested that the majority of natural Dis3L2 targets are non-coding RNAs such as tRNAs, snRNAs and snoRNAs (1,7,11,14,17–20). In human cell lines and mouse embryonic stem cells these include miRNAs such as *pre-let-7* as well as other non-coding RNAs (e.g. *Rmrp*) which have been shown to be polyuridylated and then degraded by Dis3L2 (11,18). Recent work in human cells has demonstrated a role for PARN in protecting miRNAs from DIS3L2 mediated decay thus implicating mature miRNAs as other DIS3L2 targets (21). Other direct targets which immunoprecipitated with *Drosophila* Dis3L2 included 5S rRNA and extended versions of *RNase MRP:RNA* (19). This study, along with others has demonstrated several roles for Dis3L2 in the removal of transcripts with incorrect processing or transcriptional termination (18,19). Finally, a recent study has identified a role for the uridylation-mediated DIS3L2 decay in the quality control pathway of non-sense mediated decay (NMD) in human cells (22). However, it is currently unclear how defective degradation of the identified transcripts can enhance cell proliferation.

Although previous studies have attempted to identify the proliferation pathways controlled by Dis3L2, many have used individual cells in culture rather than developing tissue which more accurately represents the physiological environment. A Dis3L2 knockout mouse model has demonstrated a transcriptional increase in *insulin growth factor 2* (*Igf2*) mRNA, however, no overgrowth was observed (23). Therefore, the cellular pathways linking Dis3L2 with proliferation have remained obscure. Understanding the cellular and molecular pathways controlled by Dis3L2 in a well characterised model system such as *Drosophila* is likely to shed light on the molecular basis of overgrowth syndromes as well as uncover novel means of controlling cell proliferation.

Here we show that the exoribonuclease Dis3L2 controls proliferation and tissue growth in *Drosophila* through regulating the levels of a conserved growth factor, Imaginal disc growth factor 2 (Idgf2). We first generated *dis3L2* null mutants plus isogenic controls using the CRISPR-Cas9 system. Using RNA-seq in a genome-wide approach, we compared the levels of RNA transcripts in *Drosophila* wing imaginal discs from *dis3L2* null mutants to that of isogenic controls. We show, in this naturally proliferative tissue, that Dis3L2 controls the expression of Idgf2 which is a member of the Chitinase-like proteins (CLP) family that is highly conserved between *Drosophila* and humans and combines properties of cytokines and growth factors. Idgf proteins have previously been shown to act in a paracrine manner to promote growth and proliferation in *Drosophila* cells. The human orthologues, chitinase-3-like-1 (CHI3L1), chitinase-3-like-2 (CHI3L2) and oviductal glycoprotein 1 (OVGP1) are upregulated in some inflammatory disorders and cancers. We also demonstrate that loss of DIS3L2 in human HEK-293T cells drives proliferation as a result of increased activity of the PI3-Kinase/AKT pathway. The data presented here suggest a novel, conserved mechanism whereby Dis3L2 regulates cell proliferation and tissue growth.

## Results

### Dis3L2 null mutants show widespread overgrowth

We have previously shown, using RNA interference, that knockdown of *dis3L2* in *Drosophila* wing imaginal discs results in increased cell proliferation (1). Knockdown of *dis3L2* specifically in the wing imaginal disc results in wings, and wing discs, that are significantly larger than controls due to cell proliferation (hyperplasia) rather than increase in cell size (hypertrophy). To further characterise the overgrowth phenotype we generated a *dis3L2* null mutant using CRISPR-Cas9, since a null mutant is more consistent than RNA interference. Using a guide RNA (gRNA) targeting the first common exon of the two *dis3L2* isoforms we generated a line with an 8bp mutation (Sup Fig 1A/B). Together with this mutant line, named *dis3L2^12^* we also generated an isogenic control line which went through the CRISPR process but remained unedited or was repaired correctly (Sup Fig 1A/B) providing an ideal control line which we named *dis3L2^wt^*.

To confirm the absence of Dis3L2 protein we generated an antibody to the first 198 amino acids which was specific to the *Drosophila* Dis3L2 protein. This confirmed the absence of Dis3L2 in *dis3L2^12^* flies (Sup Fig 1C). Crucially the *dis3L2^12^* flies showed wide-spread overgrowth when compared to the isogenic *dis3L2^wt^* control line (Fig 1). Using the null mutant we demonstrate that the overgrowth induced by the loss of *dis3L2* was not specific to the wing/wing imaginal disc (Fig 1A) with both the haltere and leg imaginal discs showing comparable, and significant overgrowth (Fig 1B). Consistent with imaginal disc overgrowth the adult flies also showed overgrowth being on average 10% larger than controls (Fig 1C).

**Figure 1:**
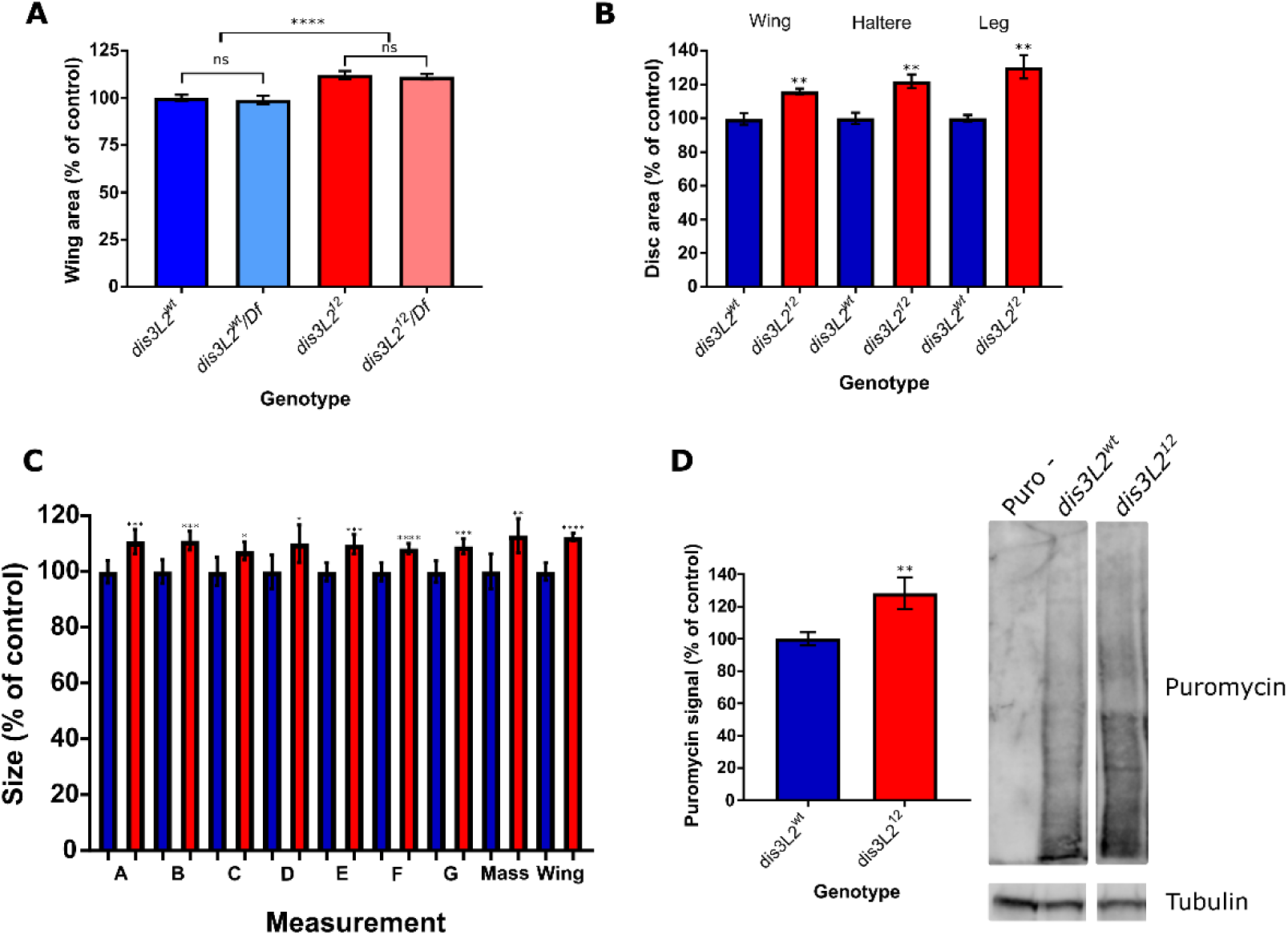
Phenotypes of *dis3L2*^12^ mutants. *dis3L2*^12^ mutants show overgrowth phenotypes. **A)** *dis3L2^12^* wings are significantly larger (14%) than control *dis3L2^wt^* wings. Hemizygous *dis3L2^12^/Df* wings are not significantly different from *dis3L2^12^* wings demonstrating that it is a null mutation. n>25, error bars represent 95% CI, **** = p<0.0001, ns = p>0.05. **B)** *dis3L2^12^* wing, haltere and leg imaginal discs are significantly larger (25%, 30%, 36% respectively) than their control counterparts. n>8, error bars represent 95% CI, ** = p<0.01. **C)** *dis3L2^12^* mutants are significantly larger than *dis3L2^wt^* flies. Calculated using measurements of various regions of the adult fly (A-G, Sup Fig 7) and total mass. n≥15, error bars represent 95% CI, * = p<0.05, ** = p<0.01, *** = p<0.001, **** = p<0.0001. **D)** SUnSET labelling performed on wing imaginal disc samples shows increased global translation in *dis3L2^12^* discs compared to *dis3L2^wt^*. Representative labelling shown along with Tubulin loading control used for quantification. Puro – represents a disc sample incubated in media only without Puromycin. n≥8, error bars represent 95% CI, ** = p=0.0057.

To genetically confirm the null nature of the allele we generated flies hemizygous for *dis3L2^12^* by crossing it over the smallest publicly available deficiency which includes the *dis3L2* locus (*Df(3L)Exel6084*). To ensure the deficiency does not affect wing area we also generated flies hemizygous for *dis3L2^wt^. dis3L2^wt^/Df* wings did not differ in area compared to *dis3L2^wt^* homozygotes showing that a single copy of wild-type *dis3L2* is sufficient to achieve correct wing area (Fig 1A) consistent with the autosomal recessive nature of Perlman syndrome. In contrast, hemizygote *dis3L2^12^* mutant wings (*dis3L2^12^/Df)* were indeed significantly larger than their respective controls confirming that this allele is null whilst also showing the specificity of the phenotype (Fig 1A). Using SUnSET labelling it is also clear that *dis3L2^12^* wing imaginal discs also show an increase in global translation (Fig 1D) which may be driving, or facilitating the increased growth of the tissue.

In addition to the overgrowth phenotype we also confirmed the fertility defect observed by Lin and colleagues (17) (Sup Fig 1D). We observed complete male infertility in both the *dis3L2^12^* homozygous and hemizygous flies. The seminal vesicles, the storage vessel of *Drosophila* sperm, were devoid of sperm supporting previous data that *dis3L2* is required for male fertility. To confirm the specificity of this phenotype we also assessed the fertility of male flies that had a ubiquitous knockdown of *dis3L2*. Using *tubulin-GAL4* to knockdown *dis3L2* throughout the fly, including the testes, results in complete male infertility demonstrating the specificity of the infertility phenotype (Sup Fig 1D). In addition to the above phenotypes, whilst viable, *dis3L2^12^* male and female adult flies show a decreased lifespan compared to *dis3L2^wt^* flies (Sup Fig 1E).

### The catalytic activity of Dis3L2 is required to control developmental proliferation

To prove that loss of Dis3L2 is indeed driving the overgrowth phenotype we first generated a number of transgenic lines to allow specific genetic rescue. There are two annotated isoforms of Dis3L2 which differ through the use of an alternative start codon resulting in two protein produces of 1032 amino acids (isoform PA) and 1044 amino acids (isoform PC). Ubiquitous expression of these lines using *actin5C-GAL4* allowed re-expression of Dis3L2 protein throughout the fly to a level consistent with that in the control line (Fig 2A). Expression of both Dis3L2 isoforms (PC) or only PA specifically in the cells fated to form the wing (using *nubbin-GAL4*) results in a complete rescue of wing area to that of *dis3L2^wt^* wings. These phenotypic rescues show that the loss of Dis3L2 is indeed responsible for the overgrowth phenotype and that isoform PA is the major isoform in *Drosophila* to regulate tissue growth.

**Figure 2:**
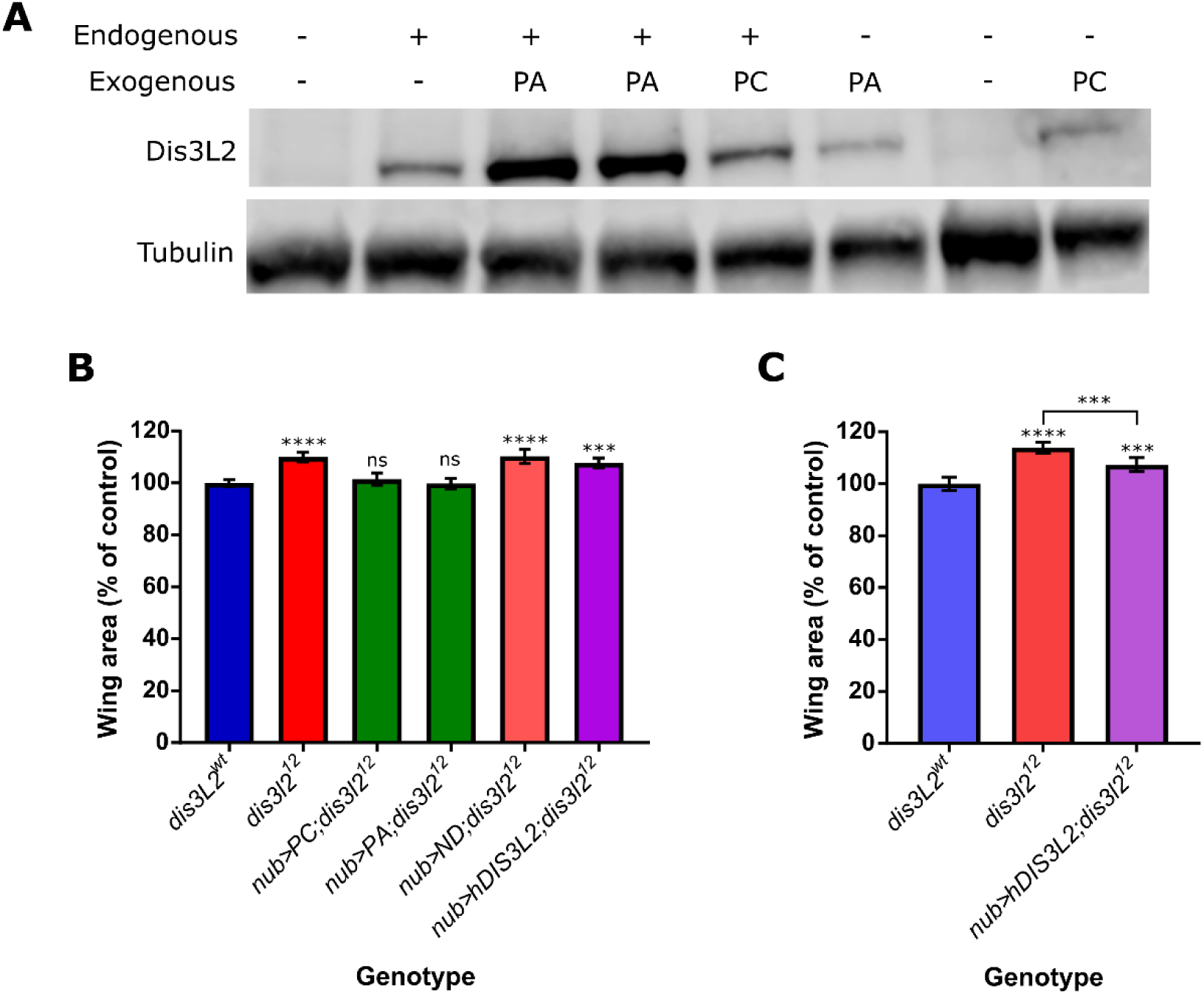
Ectopic *dis3L2* expression rescues the wing overgrowth phenotype. **A)** *UAS-dis3L2*^PA^ or *UAS-dis3L2*^PC^ can be used to express Dis3L2 in *dis3L2^12^* animals. UAS constructs were driven in either the *dis3L2^wt^* or *dis3L2^12^* animals with the ubiquitous driver *act-GAL4* with Western blots performed on whole female flies. **B)** Expression of either both Dis3L2 isoforms (PC) or just the shorter isoform (PA) in the wing pouch of the wing imaginal disc is sufficient to completely rescue wing area in *dis3L2^12^* mutants. Expression of a catalytic dead Dis3L2 (ND) does not rescue the phenotype, whilst expression of human DIS3L2 also shows no significant rescue at 25°C. n>10, error bars represent 95% CI, statistics shown refer to a comparison between the test genotype and control ***= p<0.0001, ***=p<0.001, ns=p>0.05. **C)** When driven at 29°C the human DIS3L2 construct shows a partial (~50%) rescue of wing area. n>15, error bars represent 95% CI, **** = p<0.0001, *** = p<0.001.

Next, we used a nuclease dead line (*UAS-dis3L2^ND^*) carrying a single amino acid substitution (D580N) which has previously been shown to abolish Dis3L2 catalytic activity (19), to assess the requirement of this activity on developmental proliferation. Unlike the previous rescue experiments specific expression of *dis3L2^ND^* results in no change in *dis3L2^12^* wing area demonstrating the catalytic activity of Dis3L2 is indeed essential to control proliferation in the wing imaginal disc (Fig 2B).

Given the overgrowth conditions associated with loss of DIS3L2 in humans we assessed functional homology by expressing the human DIS3L2 protein (Ensembl isoform 202, 885aa) in *dis3L2^12^* flies. We first assessed the ability of the flies to produce the human protein by driving the *UAS-hDIS3L2* construct ubiquitously with *tubulin-GAL4*. Western blotting of the resulting female progeny confirmed the presence of the human protein (99kDa) which was not due to a cross reaction of the *Drosophila* protein (118kDa) (Sup Fig 2). Expression of hDIS3L2 in *dis3L2^12^* wing cells at 25°C resulted in no change in wing area. However, as the human protein is likely to function optimally at a higher temperature we expressed hDIS3L2 at 29°C. This resulted in a partial rescue of wing area of around 50% (Fig 2C). Although not a complete rescue, the partial rescue suggests there is at least some functional homology in terms of controlling tissue growth. Previous work has shown a direct interaction between Dis3L2 and the TUTase Tailor (17,19) which occurs through domains that are not conserved in the human protein; therefore it is possible that without this interaction domain the human DIS3L2 cannot fully complement its *Drosophila* counterpart.

### Overexpression of Dis3L2 results in a reduction in tissue size and proliferation

We next wished to determine whether the activity of Dis3L2 is limited by accessory factors such as uridylyl transferases. To test this we overexpressed Dis3L2 to find out if this overexpression might cause a reduction in proliferation and tissue size. Using a publicly available line containing a P-element insertion, with an upstream activating sequence (UAS) 8bp upstream of Dis3L2 we achieved strong overexpression of the DIS3L2 protein. Ubiquitous overexpression of Dis3L2 had no effect on viability. Using *engrailed-GAL4* to drive specific overexpression of Dis3L2 in the posterior compartment of the wing imaginal disc we saw strong overexpression of Dis3L2 and a specific reduction in the posterior area of both the wing and the wing imaginal disc (Fig 3A-C). Whilst we saw a reduction in the posterior area, the anterior area, which remains wild-type for Dis3L2 expression, showed no significant difference in size when compared to the parental controls demonstrating the specificity of the phenotype.

**Figure 3:**
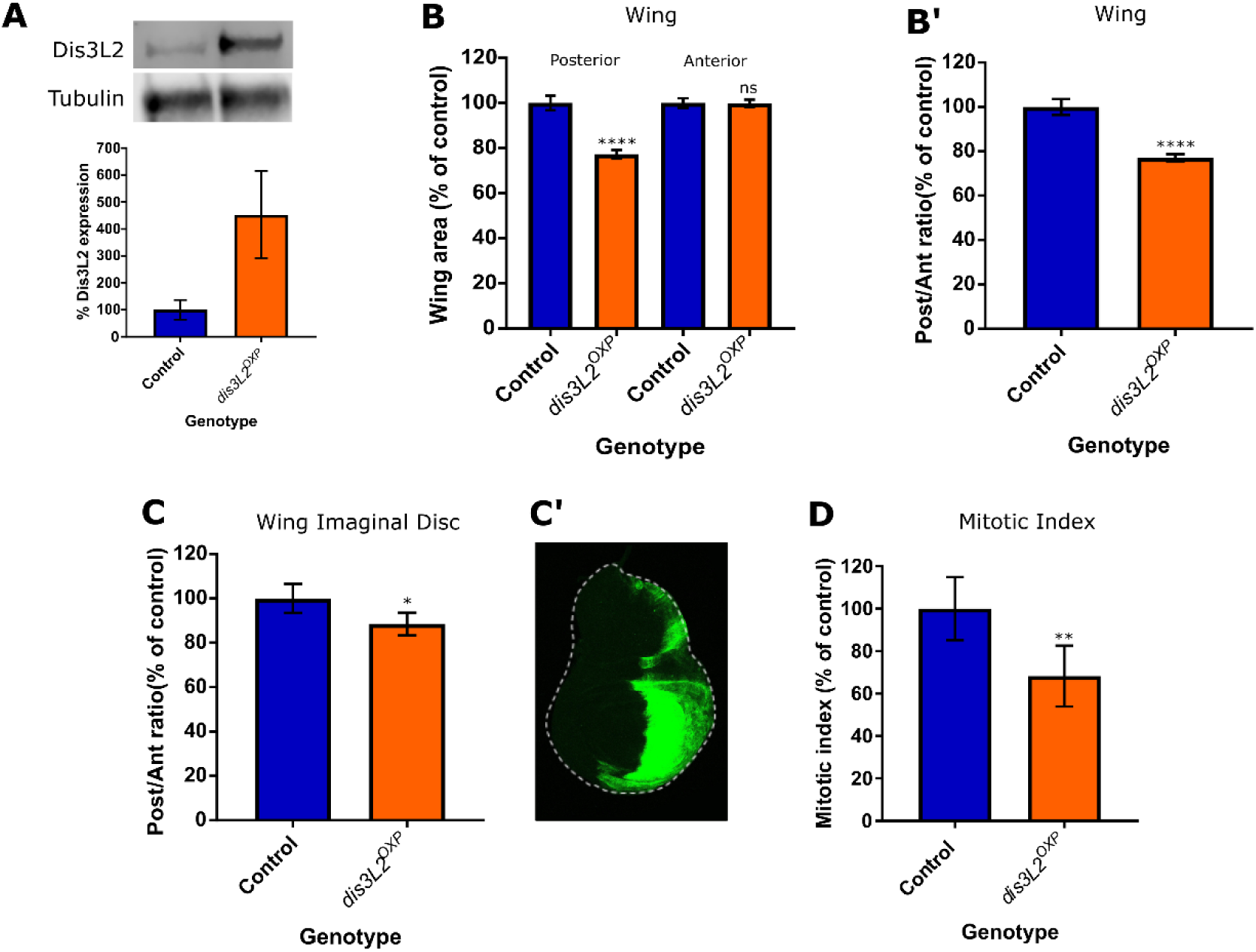
Overexpression of Dis3L2 results in a reduction in proliferation in the wing imaginal disc. **A)** Western blotting of *en-GAL4* driven Dis3L2 overexpression (*dis3L2^OXP^*) wing imaginal discs confirms a 5-fold upregulation in Dis3L2 protein throughout the disc. Control consists of the parental controls with either the *UAS* or *GAL4* insertion alone, *UAS* parent shown. n=3, error bars represent SEM. **B)** Overexpression of Dis3L2 using *en-GAL4* shows a significant reduction in the posterior area of the wing. Controls consist of parental controls to account for genetic background. n>26, error bars represent 95% CI, **** = p<0.0001, ns =p>0.05. **B’)** Normalisation of posterior area of the wing to the anterior area shows a significant reduction in the posterior area of the wing. n>26, error bars represent 95% CI, **** = p<0.0001. **C)** Overexpression of Dis3L2 in the posterior compartment of the wing imaginal disc (using *en-GAL4*) results in a reduction of the posterior area, represented as the posterior area normalised to the size of the anterior area of the tissue. The control group consists of; *en-GAL4, UAS-GFPactin/+*; to allow measurement and ensure that the *en-GAL4* insertion and expression of GFP is taken into account. n>8, error bars represent 95% CI, * = p=0.0165. **C’)** An L3 wing imaginal disc expressing *UAS-GFP* driven by *en-GAL4* to mark the posterior boundary used in **C**. **D)** Overexpression of Dis3L2 throughout the wing imaginal disc (using *69B-GAL4*) results in a reduction in the mitotic index of the wing imaginal disc. Control represents a combination of both parental controls (;;*69B-GAL4* and *;;UAS-Dis3L2*). n≥6 error bars represent 95% CI, **=p=0.0076.

To assess if this reduction in tissue size was indeed caused by a reduction in proliferation we overexpressed Dis3L2 throughout the wing imaginal disc (using *69B-GAL4*), and measured the level of proliferation using Phosphohistone H3 staining. This confirmed that the overexpression of Dis3L2 does indeed result in a reduction in proliferation and as a result a reduction in tissue size (Fig 3D). Therefore, overexpression of Dis3L2 results in reduced tissue growth. Taken together these data therefore confirms that Dis3L2 is a master regulator which plays a critical role in controlling developmental proliferation and ultimately tissue size without being limited by other accessory factors.

### RNA-sequencing reveals that imaginal disc growth factors are upregulated in *dis3L2^12^* wing discs

As Dis3L2 regulates proliferation through its catalytic activity we hypothesised that it achieves this by degrading specific, growth promoting transcripts. To identify the RNAs that become misexpressed in *dis3L2^12^* wing discs we performed RNA-sequencing on 120hr old *dis3L2^12^* and control wing imaginal discs in triplicate following rRNA depletion (see materials and methods for sample details). RNA-seq reads were processed as described in the methods. In brief, the adapters were removed, the reads were aligned to Flybase genome release 6.18 with HiSat2 (24) and subsequently assembled and analysed using the Cufflinks pipeline (25). Individual replicate comparisons were then performed to minimise false positives or false negatives. Using stringent filtering criteria to minimise the false positives (see materials and methods) we identified 501 genes that showed differential expression *dis3L2^12^* mutant wing imaginal discs (Fig 4A). Of these 344 were upregulated >1.34 fold and 157 were downregulated >1.34 fold (Sup Fig 3A). The top 30 upregulated transcripts are shown in Fig 4B. Interestingly, transcripts differentially expressed in *dis3L2^12^* wing imaginal discs have shorter 3’ UTRs (446bp) than the global average (598bp) whilst the average lengths of the 5’UTR and CDS do not differ (Sup Fig 4A) which is consistent with findings in *S. pombe* (6).

**Figure 4:**
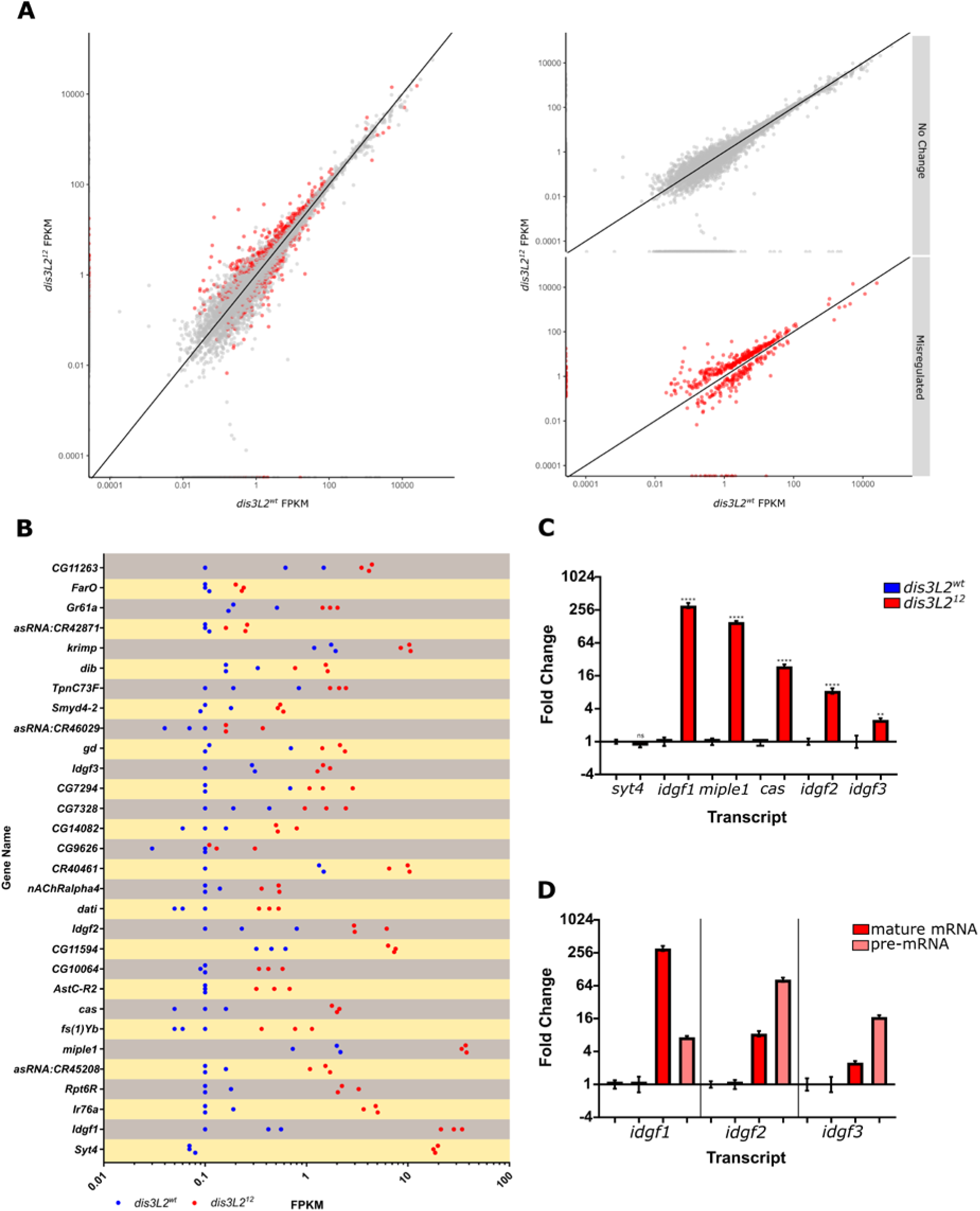
RNA-seq analysis of transcripts misregulated in *dis3L2* mutants compare to controls: **A)** 501 transcripts (red dots) were determined as differentially expressed between *dis3L2^12^* and control wing imaginal discs following stringent filtering criteria. **B)** Top 30 upregulated transcripts and their replicate FPKMs in control (blue) and *dis3L2^12^* (red) wing discs. **C)** 5 out of 6 selected transcripts showed significant increases in expression by qRT-PCR in both the sequenced samples and fresh validation samples. n=6, error bars represent SEM, ****=p<0.0001, **=p<0.01, ns=p=0.1829 using a t-test to compare the ΔΔct of *dis3L2^12^* and *dis3L2^wt^* for each transcript. D) *idgf1, 2* and *3* show at least a partial transcriptional upregulation. qRT-PCR was used to detect the pre-mRNA transcripts. n=6, error bars represent SEM, p<0.05 for all using a t-test to compare the ΔΔct of *dis3L2^12^* and *dis3L2^wt^* for each pre-mRNA.

Within those genes that showed differential expression we observed an increase in *RNaseMRP:RNA* including reads mapping to unprocessed *RNaseMRP:RNA* (Sup Fig 3B) corroborating findings in *Drosophila* S2 cells and whole flies (19) suggesting that removal of unprocessed *RNaseMRP:RNA* is a global function of Dis3L2. Gene ontology (GO) analysis of the upregulated transcripts revealed an enrichment of genes involved in carbon metabolism, symporters and glycosidases; whilst 33% of downregulated transcripts are membrane proteins (Sup Fig 3C). Interestingly, within the top 20 upregulated transcripts there are 3 imaginal disc growth factors (*idgf1/2/3*), characterised in the ‘Glycosidase’ GO term, which show 84.6, 11.8 and 7.5-fold increases respectively. In *Drosophila* there are six members of the Idgf family (Idgf1-6), however only *idgf1*, *2* and *3* appear to show Dis3L2 sensitivity as *idgf4, 5* and *6* show no changes in expression (Sup Fig 3D) indicating specific regulation. In addition to the selected Idgfs, another mRNA encoding the conserved growth factor *miple1* increased in expression by 22.3-fold.

To ensure those transcripts determined as misexpressed were not false positives we selected 6 transcripts for validation by qRT-PCR. These included the three *idgf* transcripts and *miple1*. In addition, we selected *castor*, a highly conserved transcription factor known to regulate development and proliferation of neural cells (26,27) which showed a 22-fold increase in *dis3L2^12^* tissues. Finally, we also selected *syt4* which showed the largest increase in expression (243-fold). qRT-PCR confirmed that all by *syt4* showed significant upregulation in both the samples sent for sequencing and in three additional replicates of *dis3L2^12^* and *dis3L2^wt^* (Fig 4C). To confirm the upregulation was specific and not caused by an off-target effect of the CRISPR we assessed the levels of the 5 upregulated transcripts (excluding *syt4*) in *dis3L2^12^* hemizygote wing discs where we also saw significant upregulation (Sup Fig 4B).

This observed upregulation could be a result of post-transcriptional regulation by Dis3L2, or it could be a result of indirect upregulation as a result of increased transcription. To assess this, we used TaqMan qRT-PCR assays which specifically targeted the pre-mRNAs as a transcriptional read out. The levels of pre-mRNAs were compared to the levels of the corresponding mRNAs. *idgf2 and idgf3* showed increases at the transcriptional level to an extent similar or above the increase observed for the mature transcript (Fig 4D). On the other hand, *idgf1* only showed a modest (4-fold) transcriptional increase compared to the 300-fold increase observed for the mature mRNA suggesting this was a largely post-transcriptional increase and indicating that it may be directly targeted for Dis3L2 mediated decay.

### Idgfs appear to be globally sensitive to Dis3L2 activity

Recent studies in *Drosophila* have used different approaches to assess and identify Dis3L2 targets in an embryonic cell line (S2 cells) (19) and the testes (17). To determine if the misregulated transcripts identified in our study were consistent with those observed in other studies, and whether they are tissue specific, we compared our data to that of previous studies. The publicly available raw sequencing files of RNA-seq in *dis3L2* mutant testes published by Lin and colleagues were put through our analysis pipeline (see materials and methods) to allow for direct comparison. This showed a larger number of misregulated transcripts with 1171 showing increases and 1603 showing decreases in expression.

Of the 501 transcripts misregulated in *dis3L2^12^* wing imaginal discs, 10% showed the same direction of change in the *dis3L2* mutant testes (Sup Fig 4C). Within these all three of the *idgf* mRNAs observed in our data set are also significantly upregulated in *dis3L2* mutant testes suggesting a global sensitivity of these transcripts towards Dis3L2 levels. Of the 39 transcripts that are upregulated in both *dis3L2* mutant wings and testes, 20 are also pulled down with a catalytic dead Dis3L2 in *Drosophila* S2 cells (19), including the 3 *idgfs*, demonstrating that they are physically associated with Dis3L2 suggesting at least an element of direct regulation. It is therefore possible that the transcriptional upregulation previously observed for *idgf2* and *idgf3* (Fig 4D) may be a result of feed-forward regulation. The comparative analysis gives strong evidence that, amongst others, *idgf1/2/3* are sensitive to Dis3L2 activity in a variety of tissue/cell types.

### Upregulation of *idgf2* in *dis3L2^12^* mutants drives wing overgrowth

Idgf1 and Idgf2 have previously been shown to be sufficient to drive proliferation in *Drosophila* wing imaginal disc cells. Therefore, to determine if any one of the three upregulated *idgfs* was solely responsible for driving tissue overgrowth in *dis3L2^12^* flies we knocked down each one individually in the *dis3L2^12^* mutant background. First, we assessed the efficacy of the *UAS-idgf^RNAi^* lines by driving them ubiquitously throughout the fly using *tubulin-GAL4*. Knockdown of >90% at the RNA level was confirmed by qRT-PCR (Sup Fig 5A) for each *idgf* transcript with this ubiquitous knockdown showing no obvious phenotypic defects.

To assess the contribution of the Idgfs to the *dis3L2^12^* mediated overgrowth we again used the *en-GAL4* driver to knockdown each *idgf* in the posterior compartment of the wing/wing imaginal disc allowing the anterior area as an internal control. Knockdown of either *idgf1* or *idgf3* individually within the *dis3L2^12^* mutants had no effect on wing area (Fig 5A). However, strikingly, knockdown of *idgf2* in the mutant tissues resulted in a specific reduction in the posterior area of the wing resulting in a wing area not significantly different to rescued or *dis3L2^wt^* wings (Fig 5A). Importantly, knockdown of *idgf2* in a *dis3L2^wt^* background has no effect on the posterior area (Fig 5B) showing the specificity of the role of Idgf2 in driving Dis3L2 induced overgrowth. The role of Dis3L2 in regulating Idgf2 was further shown in that re-expression of Dis3L2 also results in a reduction of *idgf2* mRNA (Sup Fig 5B). In addition levels of *idgf2* mRNA were also significantly higher in wing imaginal discs dissected from a publicly available independent line carrying a CRISPR engineered catalytic dead mutation (19) (Sup Fig 5C).

**Figure 5:**
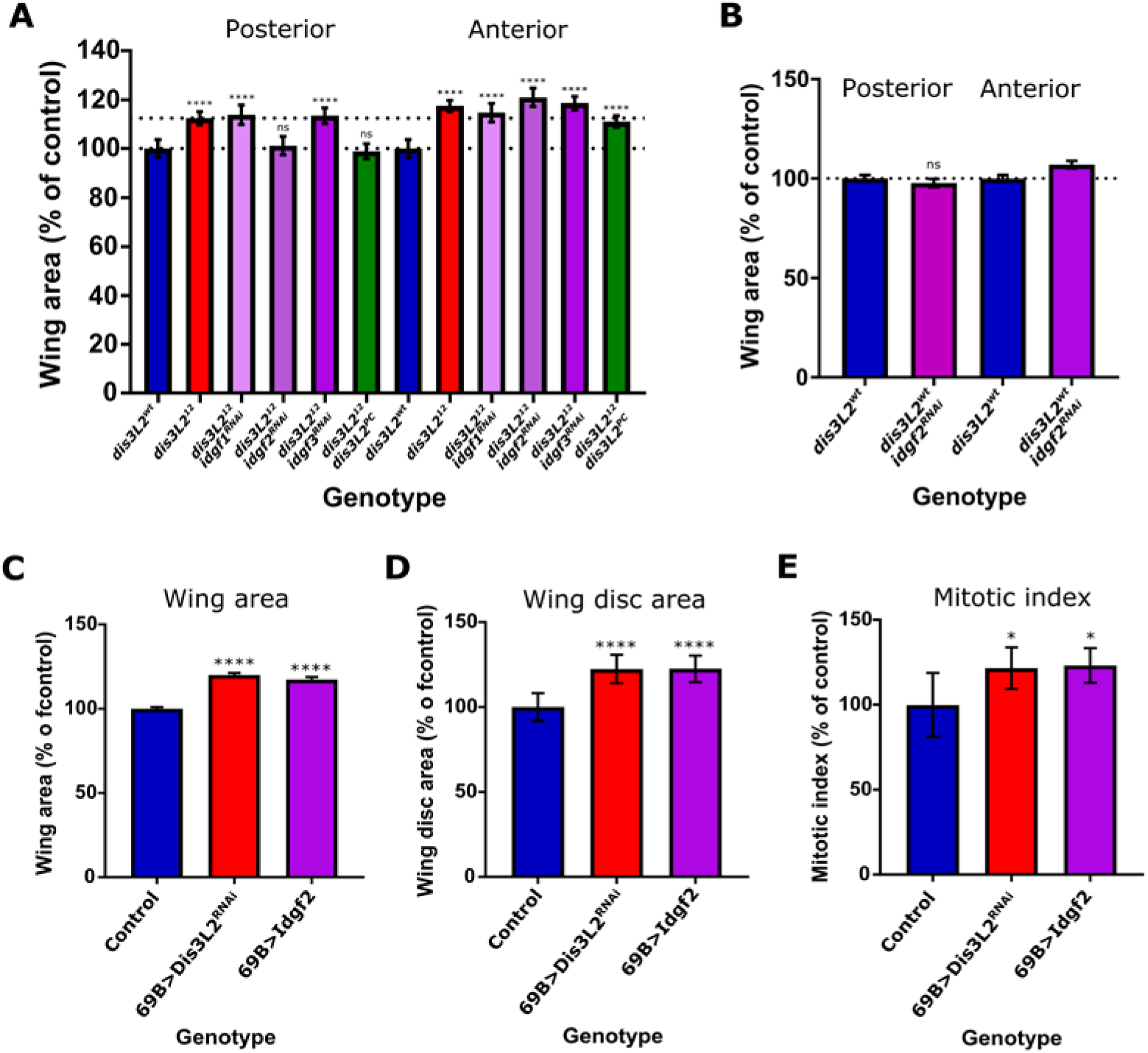
**A)** Knockdown of *idgf2* but not *idgf1* or *idgf3* in the posterior region (*en-GAL4*) of *dis3L2^12^* wing imaginal discs rescues the wing overgrowth phenotype to the same extent as expressing Dis3L2 itself. **B)** Knockdown of *idgf2* in a *dis3L2^wt^* background has no effect on wing area. Dotted lines represent average of mutant and control posterior size (relative to control). n≥19, error bars represent 95% CI, ****=p<0.0001, ns=p>0.05 represent p-values calculated for posterior and anterior area separately using an ANOVA with Tukey’s multiple comparison test comparing each area to *dis3L2^wt^*. **C/D)** Overexpression of Idgf2 throughout the wing imaginal disc using 69B-GAL4 results in overgrowth of the wing (C) and wing imaginal disc (D) to the same extent as observed when *dis3L2* is knocked down using the same method. n≥10, 95% CI, ****=p<0.0001. **E)** Idgf2 overexpression wing imaginal discs (*UAS-Idgf2/+; 69B-GAL4/+*) show increased proliferation to a level consistent with that observed in *dis3L2* deficient wing imaginal discs (*UAS-dis3L2^RNAi^/+; 69B-GAL4/+*). n≥10, 95% CI, *=p<0.05.

To ensure overexpression of Idgf2 was sufficient to drive overgrowth, we created a *UAS-Idgf2* line which increased *idgf2* expression throughout the wing imaginal disc when driven by *69B-GAL4* (Sup Fig 5D). This resulted in wing and wing imaginal disc overgrowth (Figure 5C/D) to a level not significantly different to driving *dis3L2* knockdown using the same *69B-GAL4* driver. Importantly, similarly to the loss of *dis3L2*, overexpression of Idgf2 results in overgrowth but does not disrupt tissue patterning. Finally, these Idgf2 overexpression tissues demonstrated an increase in mitotic index showing that increased expression of Idgf2 drives over-proliferation in the context of a developing tissue (Figure 5E). These data therefore demonstrate that the loss of Dis3L2 results in *idgf2* accumulation in the wing imaginal disc which in turn is sufficient and responsible for the overgrowth observed in *dis3L2^12^* animals

### Loss of DIS3L2 in HEK-293T cells results in increased proliferation

Given the lack of knowledge about the Idgf2 induced signalling pathway in *Drosophila* we turned to using human cell culture in an attempt to identify the signalling pathway activated following the loss of DIS3L2. Previous work in HeLa cells demonstrated that reduction in DIS3L2 levels results in an increase in cell number (2). Given the association between DIS3L2 and the kidney tumour, Wilms’ Tumour, we asked if the loss of DIS3L2 was sufficient to promote proliferation in HEK-293T cells, a more physiologically relevant, embryonic kidney derived cell line. Using siRNA against *DIS3L2* we achieved a maximal knockdown of 91.8% (72hrs post transfection) with a knockdown of >85% between 48 and 96 hours post transfection (144hrs post transfection a 62% knockdown was still present) (Sup Fig 6A). During this period of knockdown we observed an increase in cell number in the *DIS3L2* knockdown cells compared to a scrambled siRNA control, showing a conserved role for controlling proliferation in a clinically relevant cell type (Fig 6A).

**Figure 6:**
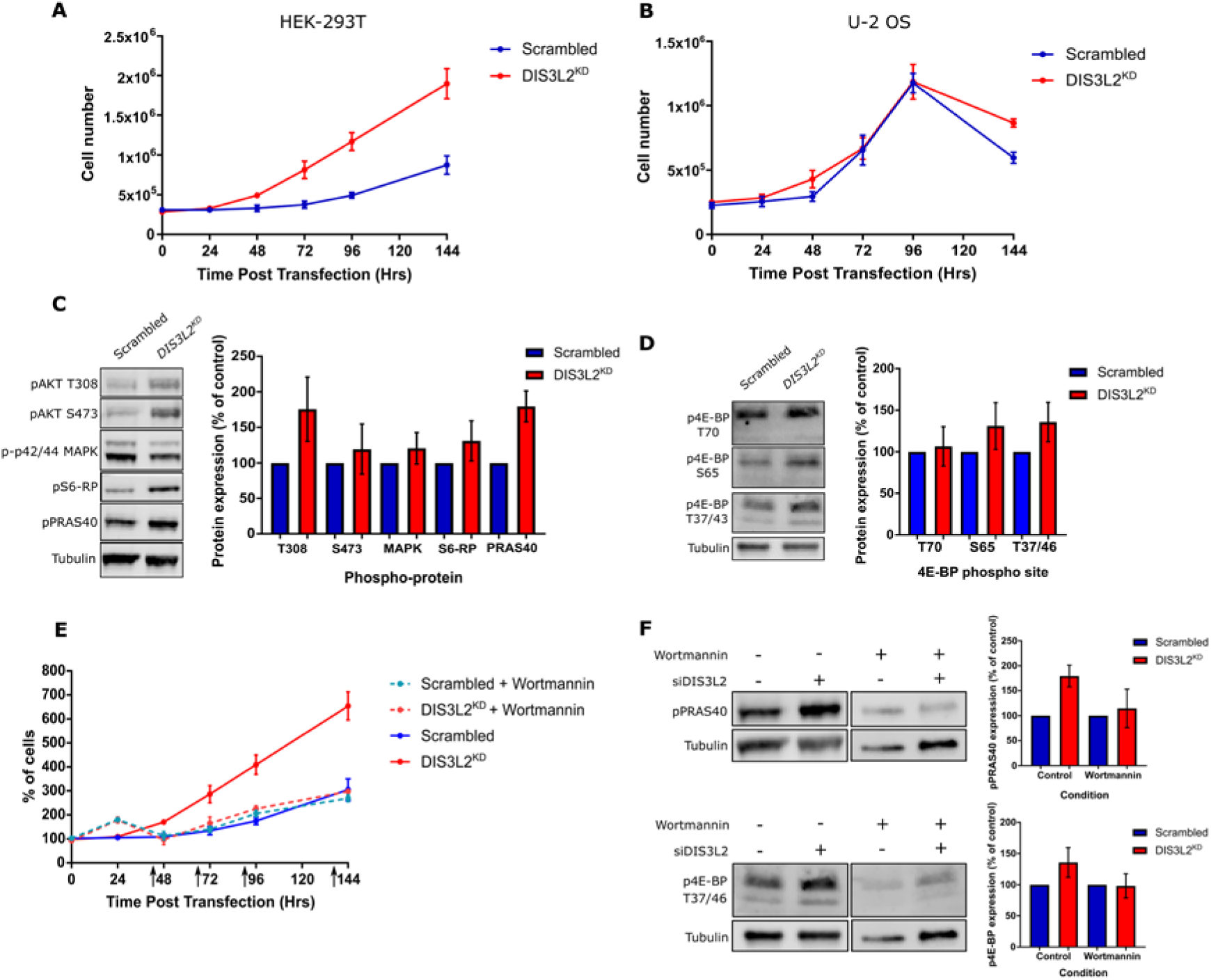
DIS3L2 depletion in HEK-293T cells results in PI3-K induced hyperplasia. **A)** Knockdown of DIS3L2 in HEK-293T cells confers a growth advantage compared to a scrambled control. n=4, error bars represent SEM. **B)** Knockdown of DIS3L2 in U2OS cells has no effect on cell number compared to a scrambled control. n=4, error bars represent SEM. **C)** Representative image and quantification of key growth promoting phospho-proteins in HEK-293T cells treated with siDIS3L2 (DIS3L2^KD^) or siScrambled (Scrambled). n=5, error bars represent SEM. **D)** Representative image and quantification of three different phosphorylation sites on 4E-BP HEK-293T cells treated with siDIS3L2 (DIS3L2^KD^) or siScrambled (Scrambled). n=3, error bars represent SEM. **E)** Treatment of DIS3L2^KD^ HEK-293T cells with Wortmannin (250nM) results in a rescue of the proliferation phenotype to that of a scrambled control with or without Wortmannin treatment. n≥3, error bars represent SEM. **F)** Western blot of pPRAS40 or p4E-BP (T37/46) with or without Wortmannin treatment in Scrambled or DIS3L2^KD^ HEK-293T cells. Quantification shows Wortmannin treatment removes the DIS3L2^KD^-induced increase in phosphorylation. n=3, error bars represent SEM.

To ask if DIS3L2-induced hyperplasia has an element of tissue specificity we knocked down DIS3L2 in the commonly used osteosarcoma cell line U-2 OS. As with the HEK-293T cells we achieved a knockdown of >90% between 72 and 144hrs post transfection (Sup Fig 6B). However, unlike HEK-293T cells, knockdown of DIS3L2 in U-2 OS cells had no effect on cell number compared to a scrambled control (Fig 6B) showing that the role for DIS3L2 in regulating proliferation does indeed have some tissue specificity in human cells. This specificity may also be a result of the developmental origin of the cells, for instance HEK-293T cells are embryonic whilst U-2 OS are from an adolescent. The conditions associated with *DIS3L2* mutations are also developmental and early onset therefore the embryonic cells may be more susceptible to DIS3L2 induced overgrowth. One may also take into account that U-2 OS cells are already cancerous, therefore may no longer be susceptible to the effects of *DIS3L2* loss.

### Loss of DIS3L2 stimulates PI3-K/AKT signalling in HEK-293T cells

In an attempt to identify the conserved molecular pathway that becomes stimulated following loss of DIS3L2 we performed a screen assigning the phosphorylation status of a number of key regulatory proteins. To assess which pathways to probe we re-analysed RNA-seq data from HEK-293T cells carrying a DIS3L2 catalytic dead mutation (20) which identified 111 differentially expressed transcripts. Gene Ontology analysis demonstrated the only significantly enriched pathway was the PI3-K/AKT pathway (enrichment score of 1.93). This would be consistent with previous data showing ectopic expression of CHI3L1 activates the AKT pathway in a variety of human cells (28–30). To test the effect of depletion of DIS3L2 on the AKT pathway we probed with p-AKT (T308 and S473) in addition to downstream pathway members. We also assessed the phosphorylation status of p42/44 MAPK, another crucial, conserved growth mediator also shown to be activated by ectopic CHI3L1/2 expression (31–33). This demonstrated that loss of DIS3L2 results in increased phosphorylation of pAKT at T308, and a complementary increase in one of its targets, pPRAS40 (Fig 6C). T308 is known to be phosphorylated by PDK1 following PI3-K activation (34), whereas S473, which shows a partial, but inconsistent increase, is phosphorylated by mTORC2 (35).

Having identified an apparent activation of PI3-K/AKT signalling in DIS3L2 deficient cells we asked if mTORC1 was activated, as might be expected for PI3-K dependent growth. We assessed the phosphorylation status of the mTORC1 target, 4E-BP on 4 conserved sites. This approach demonstrated that increased phosphorylation on S65 and T37/46 was observed in DIS3L2 knockdown cells (Fig 6D). However, we could not confirm increased T70 phosphorylation due to variation within replicates. Taken together this data suggests that DIS3L2 normally functions to inhibit the PI3-K pathway.

To phenotypically assess if the PI3-K pathway is responsible for driving overgrowth in DIS3L2 knockdown HEK-293T cells we treated them with the PI3-K inhibitor Wortmannin. Treatment of DIS3L2 knockdown HEK-293T cells with 250nM Wortmannin resulted in a complete rescue of overgrowth, demonstrating a growth prolife similar to that of the scrambled control with or without Wortmannin (Fig 6E). Wortmannin activity was confirmed by assessing the levels of pPRAS40 and p4E-BP (T37/46) by western blot which showed a reduction in phosphorylation of both proteins (Sup Fig 6C) and the removal of the increased phosphorylation status in DIS3L2^KD^ HEK-293T cells (Fig 6F). Taken together, this data demonstrates the mechanism through which DIS3L2 regulates proliferation in human cells, as is consistent with the overgrowth induced by the orthologues of Idgf2.

### PI3-Kinase contributes to over proliferation in *dis3L2^12^* wings

Having identified a role for PI3-K/AKT in *DIS3L2* knockdown HEK-293T cells we asked if the same pathway was mediating growth in the *Drosophila* wing imaginal discs. To address this, we fed larvae the PI3-Kinase inhibitor Wortmannin at 5µM and 10µM at various stages during their development. Whilst the presence of Wortmannin from 24, 48 or 72hrs into development had no effect on *dis3L2^wt^* wing area, *dis3L2^12^* wings were significantly smaller than those fed the negative control food (containing DMSO in place of Wortmannin) (Figure 7). Whilst the rescue of wing area was not complete, we saw a significant reduction in the overgrowth suggesting that Dis3L2-mediated control of growth occurs in a conserved manner through a PI3-Kinase signaling network.

**Figure 7:**
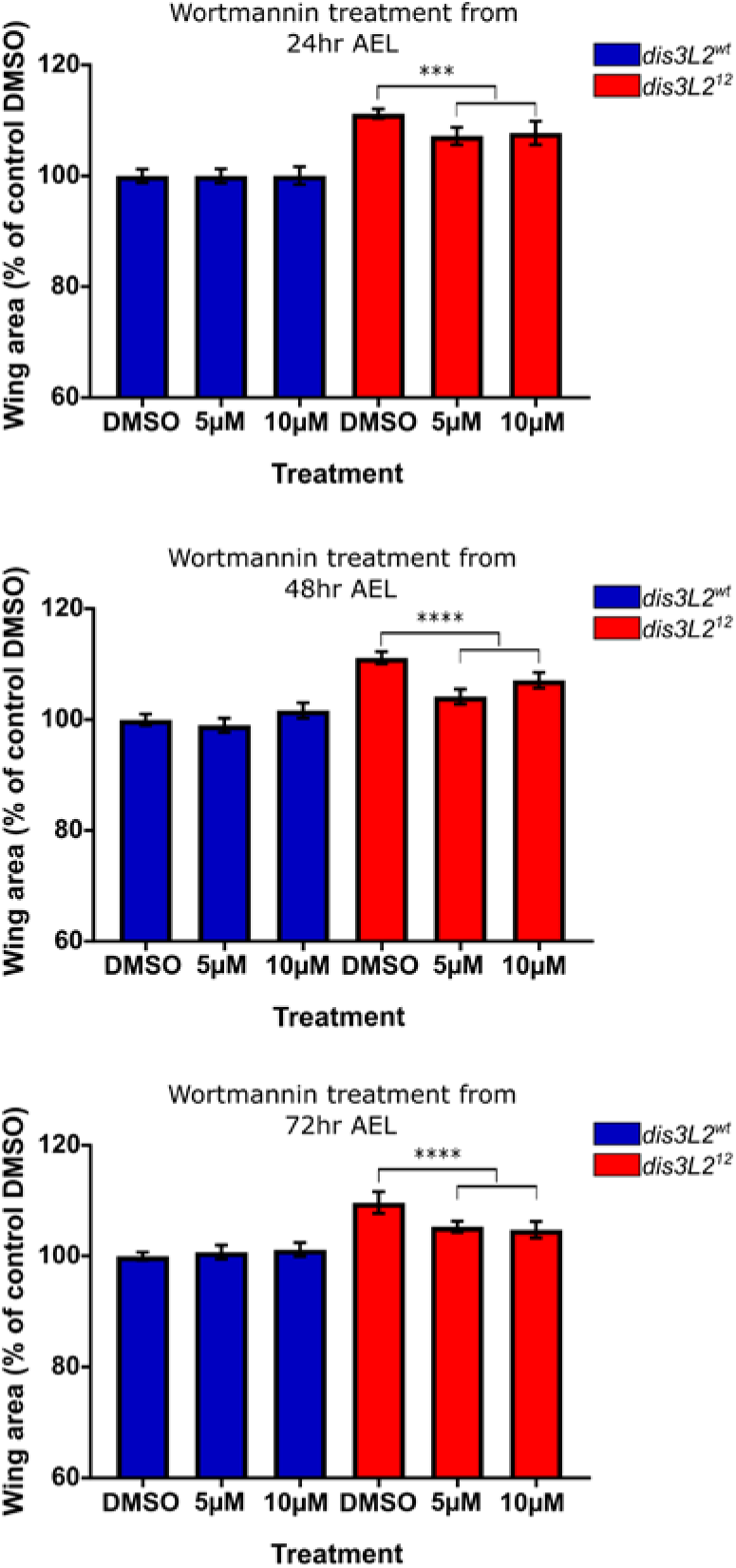
PI3-K is also involved in driving proliferation in *dis3L2^12^* mutant wings. **A)** Administration of Wortmannin at 5µM or 10µM results in a significant reduction specifically in *dis3L2^12^* wings and not *dis3L2^wt^* wings. Developing flies were cultured on Wortmannin or Control (DMSO) containing food from 24, 48 or 72 hours after egg lay (AEL). n≥16, error bars represent 95% CI, ***=p<0.001, ****p<0.0001.

Further work is required to investigate if Idgf2 directly activates the PI3-Kinase network and the identification of the receptor target of Idgf2 is crucial to this. However, here we have uncovered a conserved mechanism through which Dis3L2 elicits control over cell proliferation is tissue growth in *Drosophila* and human cells.

## Discussion

### Idgf2 co-ordinates Dis3L2 induced overgrowth in *Drosophila* partially through PI3-K signaling

In this study we have used a new *dis3L2* null mutant to identify the growth factor that is normally repressed by Dis3L2 to control tissue growth. RNA-seq analysis has revealed the loss of Dis3L2 results in the increased expression of three related imaginal disc growth factors (Idgfs), and of these Idgf2 is responsible for driving the overgrowth induced in *dis3L2* null mutant tissues. Several observations suggest that *idgf2* is a direct target of Dis3L2 in a variety of tissue and cell types. For example, we observe an increased in expression in *dis3L2* null wing imaginal discs which is rescued upon re-expression of Dis3L2. Secondly, re-analysis of RNA-seq data from *dis3L2* null mutant testes (17) also showed accumulation of *idgf1, 2* and *3*. Interestingly, this re-analysis also identified *idgf1/2/3* accumulation in *tailor* mutant testes and whilst this increase is less than that observed in *dis3L2* mutants or testes with both *tailor* and *dis3L2* mutations it suggests that *idgf2* mRNA may be targeted to Dis3L2 in a uridylation dependent manner. Crucially, re-analysis of another data set from Reimao-Pinto and colleagues also showed *idgf2* (along with *idgf1* and *3*) co-precipitating with Dis3L2 in CLIP experiments performed in *Drosophila* S2 cells (19). Finally, we also assessed the levels of *idgf2* in wing imaginal discs that have a catalytic dead mutation (19) which, like our null mutant discs, also show an accumulation of the *idgf2* mRNA. Taken together, these analyses provide strong evidence that Dis3L2 functions to control proliferation in *Drosophila* through directly degrading *idgf2* mRNA.

The requirement of Dis3L2 to actively degrade the *idgf2* is further strengthened by the data showing that whilst re-expression of wild-type Dis3L2 rescues overgrowth and *idgf2* expression, the expression of a catalytic dead Dis3L2 has no effect demonstrating the catalytic activity of Dis3L2 is essential to control tissue growth (Fig 2B). Further, we also show the potential of functional redundancy between the human and *Drosophila* proteins with a partial rescue of wing area following the expression of hDIS3L2 in *dis3L2^12^* wings at 29°C. The reasons for the lack of complete rescue could be many; for example it is possible that the human protein, which would function optimally at 37°C is not as active at 29°C. Alternatively, previous work has shown a direct interaction between the uridylyl transferase Tailor and Dis3L2 through an N-terminal coiled-coil motif, a motif which is not conserved in the human protein (17,19). Therefore, given the apparent requirement of the Tailor/Dis3L2 axis of decay in regulating *idgf2* is it possible that the lack of this domain may make the recruitment of the human protein to its targets less efficient.

Although *Drosophila* Idgf2 is the best molecularly characterised of the Idgfs in that its crystal structure has been determined (36), the downstream pathway it activates remains elusive, although data presented here suggests that Idgf2 could promote PI3-K signalling to drive proliferation. Idgf2 been shown to promote proliferation of *Drosophila* wing disc cells in culture (37,38). Further, Idgf2 was demonstrated to increase in expression by 4-fold during early stages of tissue regeneration (39). Interestingly, it has recently been shown that Idgf2 protects imaginal disc cells (C1.8) in culture from apoptosis and that this cytoprotection is associated with the induction of genes associated with energy metabolism and innate immunity (37). Our SunSET labelling, which shows increased translation in *dis3L2* mutant cells, would support the idea that these cells are more physiologically active than normal cells. It is therefore possible, that under normal conditions, Dis3L2 functions to restrain Idgf2 signalling, however, in its absence the wing disc cells are ‘primed’ in a more proliferative state to facilitate the repair of damaged tissue or protect the cells from death.

### DIS3L2 represses PI3-K/AKT signaling in HEK-293T cells

DIS3L2 loss of function mutations have been implicated Perlman Syndrome and Wilms’ tumour of the Kidney (2). To assess the role of DIS3L2 in kidney cells human cells we depleted DIS3L2 in HEK-293T cells and, as in *Drosophila*, we observed strong increases in proliferation. The role of DIS3L2 in human cells was also shown to have some specificity since the overgrowth phenotype is not observed in following DIS3L2 knockdown in the osteosarcoma cell line U2-OS. This could be for a number of reasons including the tissue of origin of the cell line, or the fact that HEK-293T cells are virally immortalised rather than cancerous. Alternatively, given that Perlman Syndrome and Wilms’ tumour are early, developmental diseases is could be that DIS3L2 is essential to maintain proliferation early in development. U2-OS cells are from an adolescent whilst HEK-293T cells are embryonic and therefore this could influence the effect of loss of DIS3L2. This would be congruent with our previous work showing that Dis3L2 is critical to control proliferation during early development (1).

Using a phospho-specific screen we observed increases in AKT/mTORC1 activity following the loss of DIS3L2 in HEK-293T cells, providing the first mechanistic explanation for DIS3L2 induced overgrowth. Interestingly, we saw increased phosphorylation of the PDK1 sensitive T308 site of AKT, however, we did not see consistent increases in S473 phosphorylation, catalyzed by mTORC2. T308 has been shown to be sufficient for AKT activity, although its activity is reduced without S473 phosphorylation (40,41). Whilst the lack of S473 activation may be technical, it may also explain the controlled tissue growth we see in *Drosophila*. For example, the partial activation of AKT would be sufficient to drive increased growth, however, without S473 activation some control over proliferation is retained. The lack of S473 phosphorylation in our experiments is also consistent with the lack of activation of the corresponding site in *Drosophila* (S505) following ectopic Idgf2 expression in C1.8 cells (37).

The contribution of the PI3-K/AKT pathway to DIS3L2 induced overgrowth was confirmed by the fact that the increased proliferation was inhibited following Wortmannin treatment DIS3L2 deficient cells. These findings are also congruent with our re-analysis of RNA-seq data from DIS3L2^ND^ HEK-293T cells (20) which shows the major differentially expressed pathway to be the PI3-K/AKT pathway. The role for PI3-K in mediating Dis3L2-induced overgrowth has subsequently been demonstrated in *Drosophila* indicating conservation of the mechanism. Whether Idgf2 directly activates this pathway, or there is co-operation with other pathways in *Drosophila* remains under investigation.

### Idgf2 is a conserved member of the chitinase-like family with orthologues implicated in a variety of pathological conditions

The importance of CLPs such as Idgf2 is shown through their extensive conservation and their association with a number of human diseases. Idgf2 is orthologous to the human CLPs CHI3L1 and CHI3L2. Elevated levels of CHI3L1 and CHI3L2 have been shown to drive proliferation of a number of human cells (28,30,32,33,42) and are associated with numerous chronic inflammatory diseases (e.g. arthritis and asthma (43–45) and poor prognosis of several cancers including glioblastoma, non-small-cell lung cancer and colorectal cancer (46–50). Interestingly, in the case of colorectal cancer, downregulation of DIS3L2 (as a consequence of knockdown of lncRNA AC105461.1) enhances the proliferation and stem-cell like properties of cells of the colorectal cancer line SW480 (51).

Strikingly, the human orthologue of Idgf2, CHI3L1 has been shown to stimulate AKT phosphorylation in a variety of cell types (28–30,42,52). Like Idgf2, ectopic expression of CHI3L1 results in an increase in proliferation in a number of human cell lines with one study showing this proliferation was PI3-K dependent (30), which is consistent with our data showing loss of DIS3L2 results in Wortmannin-sensitive overgrowth. Similarly, additional work has shown CHI3L1 protects U87 cells from death, a role again driven by PI3-K dependent activation of AKT (28). However, CHI3L1 treatment has also been shown to result in increases in MAPK phosphorylation (30,32,33) and whilst a minor increase is observed in our DIS3L2^KD^ HEK-293T cells it is neither consistent, nor significant. Consequently, it is possible that in HEK-293T cells CHI3L1 does not activate MAPK phosphorylation to a level detectable under the conditions used. Alternatively, it is possible that the activation of the AKT pathway in our cells is not driven by the Idgf2 homologue, CHI3L1 but instead is driven by another DIS3L2-sensitive growth mediator. CHI3L1 has been shown to activate the AKT and MAPK pathways by signaling through IL-13Rα and TMEM219 (29,53). Whilst IL-13Rα is not conserved in *Drosophila* it may give clues to the target of Idgf2 and it is also possible that loss of Dis3L2 in *Drosophila* also results in increased AKT signalling resulting in the overgrowth phenotypes. Although our data suggests a mechanism whereby Dis3L2 activity is critical to maintain control over CLP expression further work is clearly required to answer these remaining questions.

### How is Dis3L2 recruited to its RNA targets?

What are the molecular mechanisms whereby Dis3L2 specifically regulates the expression of *idgf2* and its other targets? Firstly, it is possible that structures within the mRNA itself render it particularly susceptible to degradation by Dis3L2. Previous work in human and *Drosophila* cells has identified a role for Dis3L2 in regulating RNA polymerase III transcripts (18–20) which have U-rich 3’ ends as a result of pol III terminators. Interestingly, MEME (54) analysis of the 3’ UTRs of mRNAs upregulated in *dis3L2^12^* mutant wing discs which also co-precipitate with Dis3L2 (19) show a significant enrichment of a U-rich motif (40% submitted transcripts), and a CA-rich motif (14.1% of submitted transcripts) which are not enriched in a control data set (mRNAs that do not change in expression or co-precipitate with Dis3L2 in S2 cells) (Sup Fig 4D/D’). Given the preference of Dis3L2 in degrading U-rich regions this could demonstrate an intrinsic mechanism facilitating Dis3L2-mediated decay of a specific set of mRNA substrates. Further, the control data set showed enrichment of a G-rich motif which was not identified in the list of likely Dis3L2 substrates (Sup Fig 4E). The inability of Dis3L2 to degrade G terminated RNAs (19), together with the fact that G-rich regions form stabilising G-quadruplex structures further strengthens the validity of our target dataset. Finally, transcripts with shorter 3’UTRs appear to be more sensitive to Dis3L2 activity (446nt vs 598nt; Sup Fig 4A) which is consistent with data from *S. pombe* (6), however, the reason behind this remains unknown.

Whilst the data presented here demonstrates a novel, and potentially conserved, mechanism through which Dis3L2 mediated decay of *idgf2* is required to control tissue growth, the downstream signalling pathway activated by Idgf2 remains elusive. Crucially, it appears that the involvement of the PI3-K in inducing Dis3L2-induceed overgrowth is conserved in human HEK-293T cells. Further understanding of these pathways is critical to understanding not only the *DIS3L2* regulated conditions but the many other conditions also driven by the increased expression of the CLP family of proteins. Exploring the endogenous role of CLP proteins in tissue culture cells is difficult due to their low levels of expression and therefore our model is an exciting and appropriate system to further unlock this largely unknown pathway of tissue growth and proliferation.

## Acknowledgements

This work was funded by a Brighton and Sussex Medical School studentship [WC003-11 to B.P.T] and a University of Brighton studentship [WC003-30] to A.L.P. The work was also supported by a Genes and Development Summer Studentship [G2162] to H.H. and a Biochemical Society Summer Vacation Studentship [G2431] to K.M.P. B.P.T was also financed by a Biotechnology and Biological Sciences Research Council grant (BB/P021042/1) to S.F.N and S.J.M. Work at ITQB NOVA was financially supported by: Project LISBOA-01-0145-FEDER-007660 (Microbiologia Molecular, Estrutural e Celular) funded by FEDER funds through COMPETE2020 - Programa Operacional Competitividade e Internacionalização (POCI) and by national funds through FCT - Fundação para a Ciência e a Tecnologia; project PTDC/BIA-MIC/1399/2014 to C.M.A and project PTDC/BIM-MEC/3749/2014 to S.C.V. R.G.M was financed by an FCT contract (ref. CEECIND/02065/2017); S.C.V was financed by program IF of “Fundação para a Ciência e a Tecnologia” (ref. IF/00217/2015).

The authors would like to thank Chris Jones, Oliver Rogoyski, Elisa Bernard, Jose Pueyo-Marques and Helen Stewart for helpful discussions plus critical reading of the manuscript. We would also like the thank Clare Rizzo-Singh for technical help.

## Author Contributions

B.P.T. designed and performed most of the experiments, prepared the figures and wrote much of the manuscript. A.L.P. performed and analysed most of the experiments on human cells. H.H. and K.M.P. performed some of the experiments. S.C.V and R.G.M performed the cloning and protein purification experiments to generate the new Dis3L2 antibody. C.M.A and S.J.M advised on the design and interpretation of the experiments. S.F.N. co-ordinated the study, contributed to the design and interpretation of the experiments and contributed to the writing of the manuscript.

## Methods

### Drosophila Husbandry

Fly stocks were cultivated on standard media in uncrowded conditions at 25°C unless otherwise stated. The following stocks were obtained from Bloomington Stock Center: *tub-GAL4* (*P{w^+mC^=tubP-GAL4}LL7* originally from stock 5138, *y^1^ w* ;; P{w^+mC^=tubP-GAL4}LL7/TM6b,GFP*), *act5C-GAL4* (stock 4414, *y^1^ w*; P{w^+mC^=Act5C-GAL4}25FO1/CyO, y^+^ ;*), *nub-GAL4* (stock 25754; *P{UAS-Dcr-2.D}1, w^1118^; P{GawB}nub-AC-62*), *69B-GAL4* (stock 1774; *w*;; P{GawB}69B*), *UAS-idgf1^RNAi^* (stock 57508, *y^1^sc*v^1^ ; P{TRiP.HMC04823}attP40*), *UAS-idgf2^RNAi^* (stock 55935, *y^1^sc*v^1^ ; P{TRiP.HMC04223}attP40*), *UAS-idgf3^RNAi^* (stock 67226, *y^1^sc*v^1^ ; P{TRiP.HMC06327}attP40*), *Df(3L)Exel6084* (stock 7563, *w^1118^ ;; Df(3L)Exel6084, P{XP-U}Exel6084/TM6B*). Stocks obtained from the Vienna Drosophila Resource Center were: *UAS-dis3L2^RNAi^* (stock v51854, *w^1118^ ; P{GD9240}v51854* ; and stock v100322, *;; P{KK105902}VIE-260B*) and Dis3L2^CD^ (stock 312503). *en-GAL4 (*A kind gift from Paul Martin, *; engrailed-GAL4,UAS-GFPactin/Cyo ;)* To overexpress Dis3L2 the stock *w* ;; P{GSV3}GS6090/TM6* was purchased from the Kyoto Stock Center (stock 200902), however, a mutation outside the Df6084 locus caused homozygous lethality; this was repaired by recombination.

### Generation of CRISPR mutants

A gRNA to the first exon of *dis3L2* was cloned into the Bbs1 site of pCFD3-dU6:3. Cloned vector was then sent to BestGene Inc for injection into vas-Cas9 embryos (BDSC 51323 *y^1^ M{vas-Cas9}ZH2A w^1118^*). Injected larvae were then returned, and potential mutant chromosomes were balanced and subject to PCR screening using the gRNA as a forward primer (see supplemental Table 1 for all primers/gRNA used in this study). Potential mutant stocks were sequenced at Eurofins Genomics. An 8bp frame shift mutation was identified and its lack of Dis3L2 confirmed by Western blotting. This line was named *dis3L2^12^*.

### Western blotting and generation of anti-dDis3L2 antibody

A polyclonal antibody was raised against the N-terminal tagged Dis3L2 peptide consisting of the first 198 amino acids of the *Drosophila* protein. pDNR plasmid-B527503, containing the cDNA for *Drosophila* CG16940 RC gene (Isoform PC), was used as template for the amplification of the *Drosophila dis3L2* gene in the construction of two N-terminal fusion proteins with different tags (GST-Dis3L2 and MBP-Dis3L2), that were expressed and purified in *E. coli*. The GST- and MBP-tagged proteins were used for the immunization in the goat and for the affinity purification of the anti-serum, respectively (by SICGEN, Portugal).

Western blots were performed in samples of adult females (x4), L3 larvae (x5) or wing imaginal discs (x60). Tubulin was used as a loading control on all blots. All blots were blocked in Odyssey Blocking Buffer (PBS) (LI-COR cat. no. 927-40000). Mouse anti-Tubulin primary antibody (Sigma, cat. no. T9026) was used at 1:2000 dilution. Goat anti-Dis3L2 (produced in this publication) was used at 1:2500. Anti-mouse and anti-goat fluorescent secondary antibodies were used at 1:20,000 (LI-COR Donkey anti-mouse IRDye 800CW (cat. No. 925-32212) and Donkey anti-goat IRDye 680RD (cat. no. 925-68074). Anti-human DIS3L2 (hDIS3L2) was used at 1:500 (Novus Biologicals cat. No. NBP-1-84740) with Goat anti-Rabbit IRDye 680RD secondary antibody (LI-COR cat. no. 925-68071). Human anti-GAPDH (Abcam cat. No. ab8245) was also used at 1:10,000 as a human specific loading control along with the anti-mouse IRDye 800CW secondary antibody above. Detection and quantification were performed using the LI-COR Odyssey FC imager and Image Studio (version 5.2) respectively.

### Phospho-specific western blotting in HEK-293T cells

Blots were performed on cell pellets from the specified time post transfection or treatment. Samples were ran on 4-12% Novex gradient gels (Invitrogen cat. no NP0321BOX), apart from those probing p4E-BP for which Mini-Protean TGX Stain-Free 4-15% gels were used (Bio-Rad cat. no. 4568084). All membranes were blocked in 3% BSA in TBS with antibody incubation in 3% BSA in 0.1% TBS-Tween. Primary antibodies used were as follows: Anti-pAKT (T308) (Cell Signalling cat. no. 1303S), anti-pAKT (S473) (Cell Signalling cat. no. 4060S), anti-pp42/44 MAPK (T202/Y204) (Cell Signalling cat. no. 9101S), anti-pPRAS40 (T246) (Cell Signalling cat. no. 2997S), pS6-Ribosomal Protein (Cell Signalling cat. no. 2215S), anti-p4E-BP (T37/46) (Cell Signalling cat. no 2855S), anti-p4E-BP (S65) (Cell Signalling cat. no. 9451S) and anti-p4E-BP (T70) (Cell Signalling cat. no. 5078S) were all used at 1:1000. Goat anti-Rabbit IRDye 680RD secondary antibody (LI-COR cat. No. 925-68071) was used followed by detection and quantification using the LI-COR Odyssey FC imager and Image Studio (version 5.2) respectively.

### Assessing lifespan

1 day old *dis3L2^wt^* and *dis3L2^12^* flies were transferred into new food. Every 3-4 days flies were transferred into new food and the number of deceased flies were counted. This continued until all flies had deceased.

### Measurement of wing and wing imaginal disc area

Flies were aged to between 1 and 2 days old for all measurement experiments. A single wing was cut from each fly and stored in isopropanol for 1 hour before being mounted in DPX (Sigma cat. no. 06522). All results shown are for male wings, however, female wings also showed the same phenotypes. Mounted wings were measured using Axiovision 4.7 on an Axioplan microscope (Carl Zeiss). Imaginal discs were dissected, in PBS, from L3 larvae 120hr old, aged using a 1 hour egg lay. Dissected discs were mounted on Poly-L-Lysine slides and mounted in 85% Glycerol. Disc areas were measured using Axiovision 4.7 on an Axioplan microscope (Carl Zeiss).

### Measurement of whole flies

Control and mutant female flies were left for 1 hour to lay on grape agar plates at 25°C. 24 hours later 20 L1 larvae were transferred into food vials and left to develop in uncrowded conditions. 5 vials (100 larvae total) were set up for each genotype. Eclosing adults were then aged to 1 day old, individually weighed and photographed. Specific, defined regions of each male fly were then measured using ImageJ (Sup Fig 7).

### SUnSET labelling

Surface sensing of translation (SUnSET) was performed on *dis3L2^wt^* and *dis3L2^12^* wing imaginal discs. Wing discs were dissected in batches of 30 and incubated in fully supplemented Shields and Sang M3 insect medium (Sigma-Aldrich, cat. no S8398) containing 2μg/ml Puromycin (Sigma-Aldrich, cat. no. P8833) for 1 hour at 25°C with rotation. Western blotting was performed to determine the levels of Puromycin incorporation with Tubulin as a loading control. Mouse anti-Tubulin primary antibody (Sigma-Aldrich, cat. no. T9026) was used at 1:2000 dilution. Mouse anti-Puromycin (clone 12D10) primary antibody (Merck Millipore, cat. no. MABE343) was used at 1:1000 dilution. Anti-mouse IRDye 800CW secondary antibody (LI-COR Biosciences, cat. no 926–32210) was used at 1:20 000 dilution to detect both primary antibodies. Each sample was run in parallel on two gels/membranes so the Tubulin band could be distinguished from Puromycin containing peptides. Quantification of Tubulin (50 kDa) and Puromycin peptides (from smallest size visible to 245 kDa) was achieved using LI-COR Biosciences Image Studio software.

### Fertility assays

Single virgin, 3 day old male flies of the test genotype were crossed to 2 virgin *dis3L2^wt^* females. Flies were left for 48 hours to mate, and then removed. Eclosing progeny were counted for 7 days. In all *dis3L2^12^* homozygous and hemizygous male replicates many eggs were laid but none were fertilised.

### Generation of *UAS-dis3L2* and *UAS-idgf2* lines

*Drosophila dis3L2* was amplified from pDNR-DUAL (DGRC clone BS27503) using two primer pairs; (1) PC which amplified the full coding region of *dis3L2* capable of encoding both isoforms and (2) PA which only amplified the coding region of the shorter isoform (PA). These were cloned into the AgeI and XbaI sites of pUASP-attb. Constructs were then sent to BestGene Inc for injection into a recipient line with an attP site at 51C1 (BDSC 24482; M{3xP3-RFP.attP’}ZHC51C). Lines where the insert was successfully integrated were confirmed by PCR and balanced. To produce the *UAS-dis3L2^ND^* line site-directed mutagenesis was performed using the Agilent Quickchange Lightning mutagenesis kit (Agilent, cat. No. 210518), and primers containing the G1738A mutation (D580N) on the pUASP-attB vector containing the PC insert. Human *DIS3L2* (Ensembl isoform 202) was amplified from oligo(dT) primed cDNA produced from hFOB cells (ATCC CRL-11372) and cloned into the NotI and SpeI sites of pUASP-attB. Injection and subsequent processing were performed as above. *UAS-idgf2* was produced by amplifying *idgf2* (CDS and 3’UTR) from cDNA from *dis3L2^wt^* wing imaginal discs and subsequent cloning to the NotI and SpeI sites of pUASP-attb. The created vector was then injected into attP40 (y^1^ w^67c23^ ; P{CaryP}attp40) and balanced. The sequences of all primer sets used are described in Supplemental Table 1.

### Mitotic index

Immunocytochemistry was performed essentially as described in (1) on 120hr L3 wing imaginal discs. Images were taken with a Leica SP8 confocal microscope. Anti-Phosphohistone H3 (Cell Signalling, cat no. 9701) was used at 1:300 dilution. Cy3-conjugated monoclonal Donkey anti-mouse IgG secondary antibody was used at 1:400 (Jackson ImmunoResearch, cat. no.715-165-151). The number of nuclei undergoing mitosis were counted using the ImageJ plugin DeadEasy MitoGlia (55). The mitotic index was then calculated for each disc by dividing the number of cells in M phase by the area of the disc.

### Wortmannin assay in *Drosophila*

Fresh food contaning Formula 4-24 Instant *Drosophila* Medium (Carolina cat no. 173200), yeast and bromophenol blue was made each day with either the desired concentration of Wortmannin (5µM or 10µM) or an equal volume of DMSO. 4 hour egg lays were performed on grape agar plates and an equal number of larvae were transferred to Wortmannin or control food for each genotype at specific developmental time points (24hr, 48hr and 72hr after egg lay) then left to develop at 25°C. Consumption of the drug was confirmed by visualising the blue food in the gut. Wings are then cut and measured from eclosing adults as outlined above.

### RNA-seq sample preparation and analysis

RNA was extracted from 60 wing imaginal discs dissected from 120hr L3 larvae. Three replicates of control and mutant were dissected. In these experiments, control samples were collected from a stock containing a 6bp mutation in *dis3L2* that does not affect wing size or induce any other obvious phenotype. Downstream qRT-PCR validation samples were collected from the *dis3L2^wt^* stock which contains the wild-type sequence and were grouped together with the sequenced samples for downstream analysis (*dis3L2^wt^*). To ensure accurate aging, 3 hour egg lays were used and wing discs were dissected in Ringers solution 120hrs after egg laying. RNA was extracted using a miRNeasy RNA extraction kit (Qiagen, cat. no. 217084), with on-column DNase digestion (Qiagen, cat. no. 79254). RNA concentrations were measured on a NanoDrop One spectrophotometer (Thermo Scientific). RNA integrity was assessed on an Agilent 2100 Bioanalyser.

400ng total RNA was depleted for ribosomal RNA as performed in (56) with the resulting RNA sent to Leeds Genomics for library preparation using the Illumina TruSeq Stranded protocol. Subsequent libraries were run in a paired-end sequencing run on a HiSeq 3000 generating between 36 and 44 million reads per sample. Raw RNA-sequencing files have been deposited in ArrayExpress. Accession number: E-MTAB-7451

Sequencing quality was assessed using FastQC c0.11.7 (http://www.bioinformatics.babraham.ac.uk/projects/fastqc/) and adapters were removed using Scythe v0.993b (https://github.com/vsbuffalo/scythe). Further quality control and read trimming was achieved using Sickle v1.29 (https://github.com/najoshi/sickle). The remaining high quality reads were mapped to the *Drosophila melanogaster* genome from Flybase (r6.18 (57)) using HiSat2 v2.1.0 (24). Differential expression was completed and normalised FKPM values were generated using the Cufflinks pipeline (25). Individual replicates from each condition showed good consistency (Supplemental File 1). Due to issues with statistical outputs from these pipelines we used individual replicate comparisons to minimise false positives when specifying that a gene is differentially expressed. All mutant replicates required a fold change of >1.34 compared to all control replicates. A fold change cut off of >1.34 was selected as this was the smallest fold change deemed significant from the Cuffdiff output. Alignment results and non-default parameters used during the analysis are shown in Supplemental File. Raw sequencing files from (17) were processed in the same analysis pipeline to allow direct comparison.

### RNA extraction and qRT-PCR

RNA extractions were performed using a miRNeasy RNA extraction kit (Qiagen, cat. no. 217084), with on-column DNase digestion (Qiagen, cat. no. 79254). RNA concentrations were measured on a NanoDrop One spectrophotometer (Thermo Scientific).

For qRT-PCR, 1µg of total RNA was converted to cDNA in duplicate using a High Capacity cDNA Reverse Transcription Kit (Life Technologies, cat. no. 4368814) with random primers. A control “no RT” reaction was performed in parallel to confirm that all genomic DNA had been degraded. qRT-PCR was performed on each cDNA replicate in duplicate (i.e. 4 technical replicates in all), using TaqMan Universal PCR Master Mix, No AmpErase UNG (Life Technologies, cat. no. 4324018) and an appropriate TaqMan assay (Life Technologies). For custom pre-mRNA assays, the pre-mRNA sequence of the desired target area was submitted to Life Technologies’ web-based Custom TaqMan Assay Design Tool as in (58) (Supplemental Table 1). *rpL32* (*rp49*) was used for normalisation.

### Human cell culture and growth curve analysis

HEK-293T and U2-OS cells were cultured in Dulbecco’s Modified Eagle’s Medium/F12 (DMEM/F12 – Gibco cat. no. 21331-020) supplemented with 10% foetal bovine serum (PAN Biotech, cat. no. P40-37500), 2mM L-Glutamine (Gibco cat. no. 25030-024) and antibiotics (100IU/mL penicillin, 100µg/mL streptomycin, Sigma Aldrich cat. no. 15140-122), at 37°C in a 5% CO_2_ humidified incubator. For growth analysis of DIS3L2 knockdown, 3×10^5^ cells were plated in 6 well plates and transfected with 30pmol siRNA (DIS3L2 (Invitrogen cat. no. AM16708)/scrambled (Invitrogen cat. no. AM4611)) using Lipofectamine RNAiMAX reagent (ThermoFisher cat. no. 13778030). Cell number was counted in triplicate every 24hrs for 144hrs. For Wortmannin treatment analysis cells 3×10^5^ HEK-293T cells were plated and treated with siRNA as above. 250nM Wortmannin was then added 44, 68, 92 and 140hrs post-transfection. Cells were counted in triplicate every 24hrs following transfection.

### Statistical tests

All statistical analyses were performed in either R v3.5.1 or GraphPad Prism 7. Two-sided two-sample t-tests were used to compare the means of single test groups to single control groups. If multiple comparisons were required, a one-way ANOVA was performed with a post-test to compare the means of each possible pair of samples.

## Supplemental Information

**Supplemental Figure 1:**
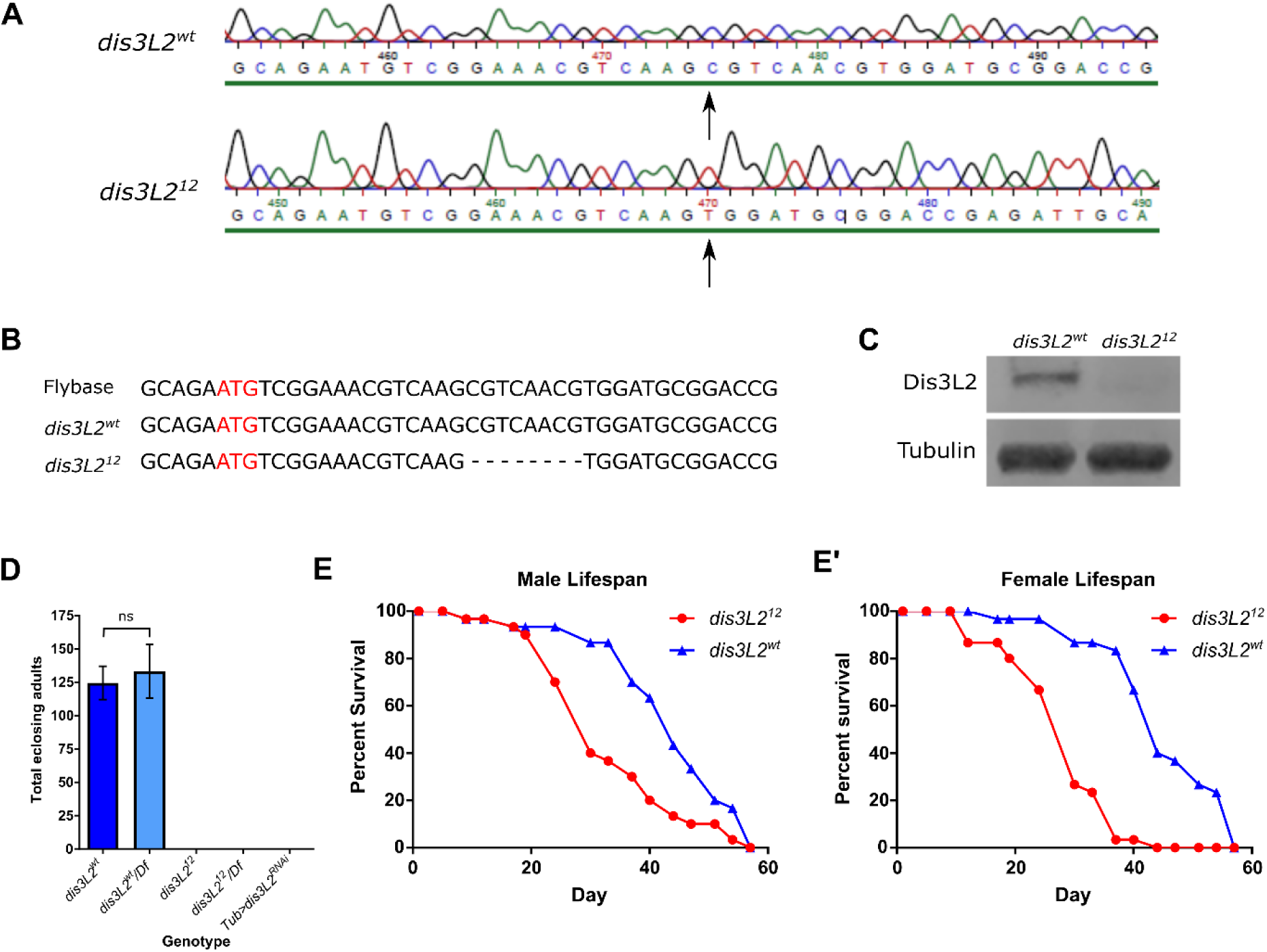
Generation and characterisation of a CRISPR-Cas9 *dis3L2* null mutant. **A)** DNA sequencing of the control *dis3L2*^wt^ and mutant *dis3L2^12^* lines. **B)** Alignment of control *dis3L2*^wt^ and mutant *dis3L2^12^* sequences together with a sequence obtained from Flybase (57) showing an 8bp mutation in *dis3L2^12^*. Start codon of isoform PA shown in red. **C)** Western Blot demonstrating an absence of Dis3L2 protein in the *dis3L2^12^* null mutant line with Tubulin used as a loading control. n>5. **D)** In all sample crosses no progeny were observed for any of the homozygous, hemizygous or knockdown mutants. n>5, error bars represent 95% CI. **E/E’)** *dis3L2^12^* mutant males **(E)** and females **(E’)** have a reduced lifespan compared to *dis3L2^wt^* isogenic controls. n≥30 flies per sex.

**Supplemental Figure 2:**
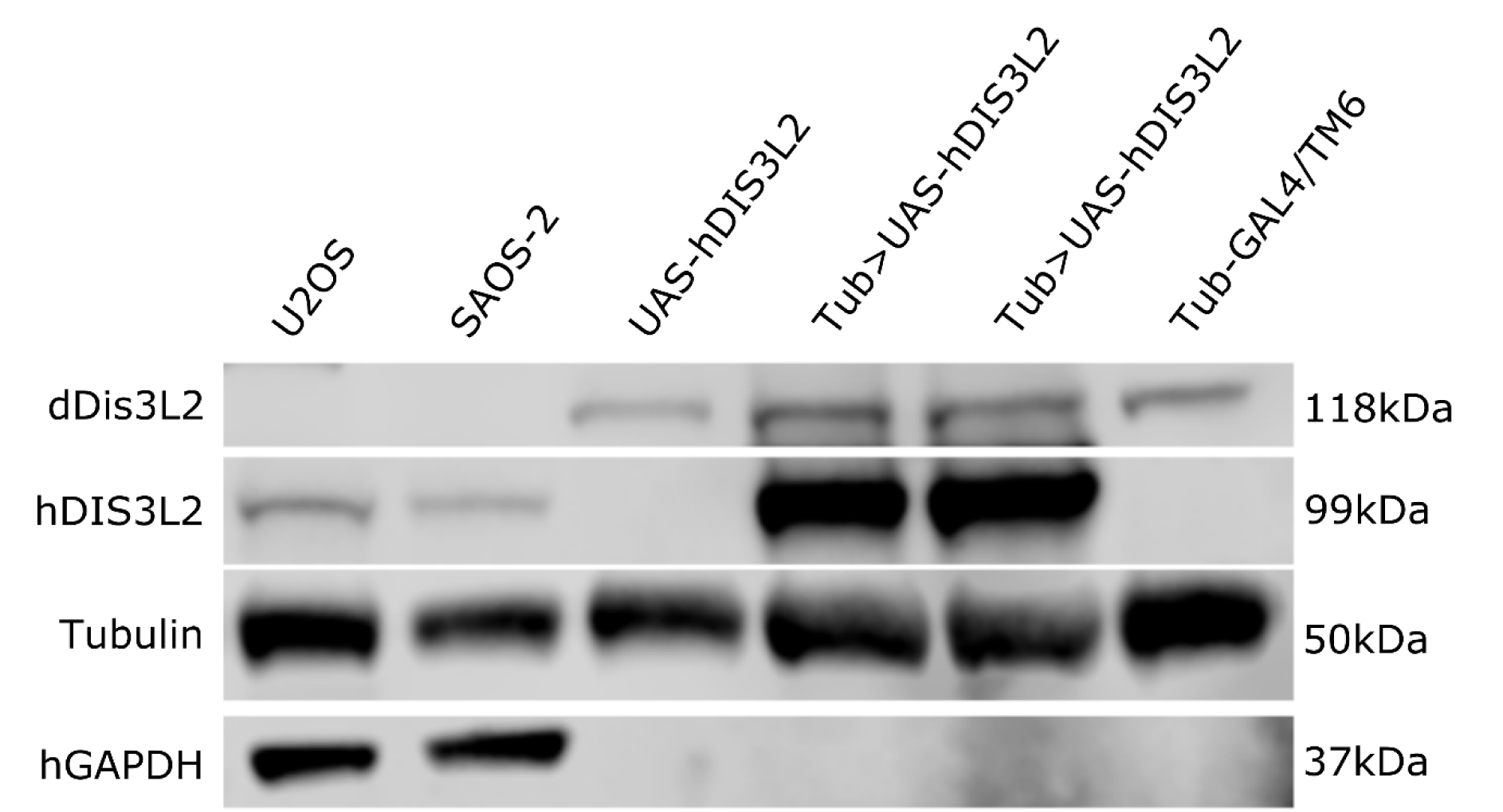
Human DIS3L2 is successfully expressed in *Drosophila* from the *UAS-hDIS3L2* construct. Two human osteosarcoma cells lines were used as positive controls (U-2 OS and SAOS-2). Protein lysate was prepared from 1×10^6^ cells or 4, 1 day old, adult females. hDIS3L2 is observed specifically in the human cells and females where *UAS-hDIS3L2* had been driven by *tub-GAL4* at 25°C (*Tub>UAS-hDIS3L2*). No product was observed in the parental controls (*UAS-hDIS3L2* and *Tub-GAL4/TM6*), confirming no ‘leaky’ expression from the *UAS-hDIS3L2* line. Tubulin used as a loading control for all samples as the antibody detects both human and *Drosophila* protein. hGAPDH used as an additional loading control specifically for the human cell line samples.

**Supplemental Figure 3:**
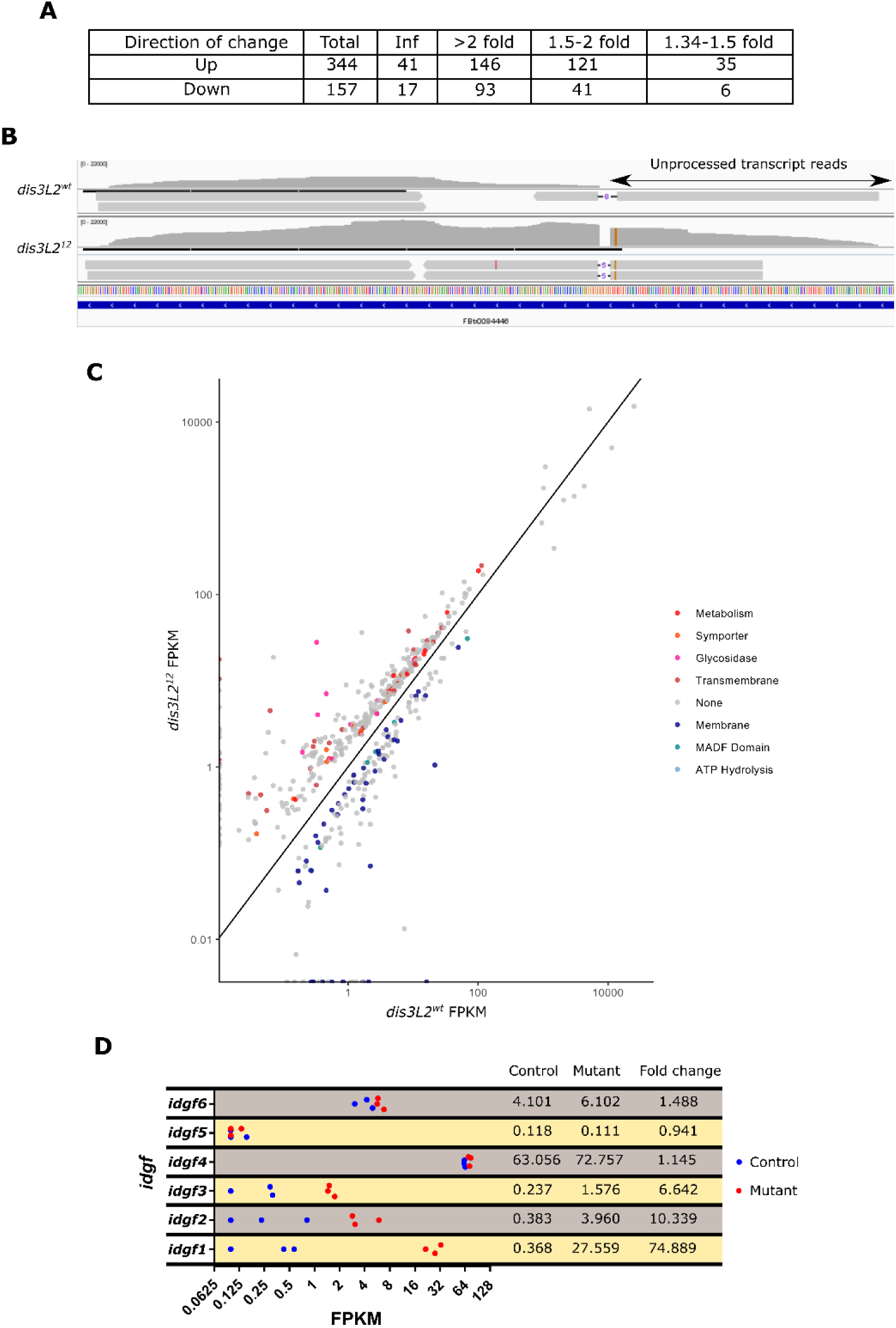
Summary and highlights of the analysis of the RNA-seq experiment. **A)** Summary of the number of transcripts showing up- and downregulation in *dis3L2^12^* wing imaginal discs. A fold change cut off of >1.34 was selected as this was the smallest change deemed significant by Cuffdiff. Inf change represents transcripts that were only detected in a single condition. **B)** Integrative Genomics Viewer screenshot showing accumulation of unprocessed *RNaseMRP:RNA* transcripts in *dis3L2^12^* tissues. **C)** Scatter plot of misregulated genes coloured by significant gene ontology categories. “None” represents genes that belonged to a category that was not significantly enriched. **D)** Strip plots showing replicate FPKM values for each of the *idgf* family in *dis3L2^wt^* and *dis3L2^12^* wing discs. Only *idgf1, idgf2* and *idgf3* show changes in expression.

**Supplemental Figure 4:**
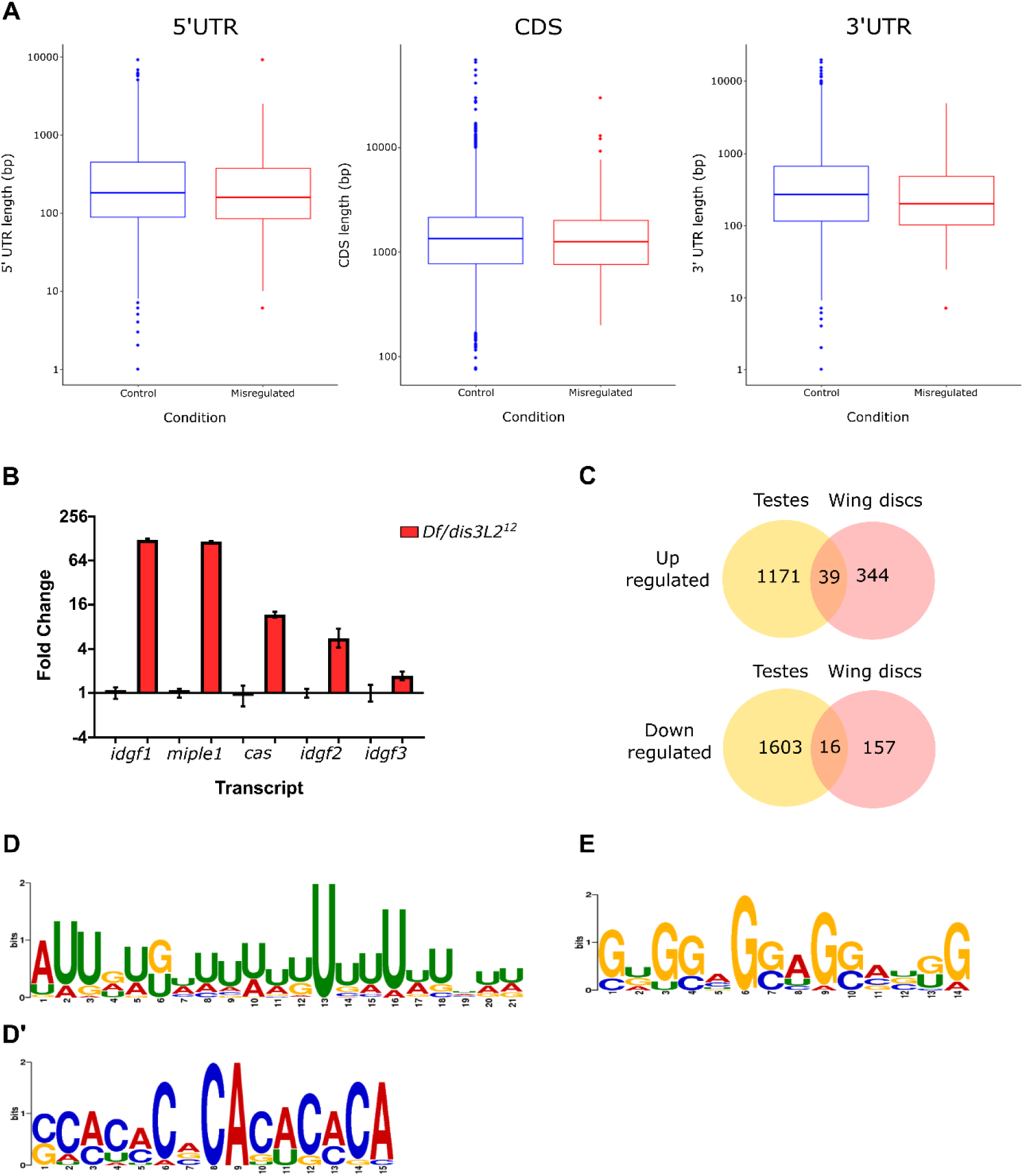
Additional information from RNA-seq data. **A)** Transcripts misexpressed in *dis3L2^12^* wing imaginal discs have shorter than average 3’ UTRs (446nt vs 598nt). 5’ UTRs (337nt vs 365nt) and the coding sequence (CDS, 1712nt vs 1837nt) show no difference in length between transcripts. Median and upper and lower quartile are represented by horizontal lines with maximum and minimum values show vertically. **B)** All validated mRNAs show significant increases in expression if *dis3L2^12^* hemizygote wing imaginal discs compared to *dis3L2*^wt^ wing discs. n≥3, error bars represent SEM, p<0.05 for all. **C)** 10% of transcripts misexpressed in *dis3L2^12^* wing discs are also misexpressed in *dis3L2* mutant testes (17). **D/D’)** MEME analysis identifies U-rich and CA-rich motifs are significantly enriched in likely Dis3L2 targets. **D)** U-rich: E-value = 5.2e^−6^, found in 40.6% of submitted sequences. **D’)** CA-rich: E-value = 0.0012, found in 14.1% of submitted sequences. **E)** A G-rich motif present in 23.1% of control sequences (sequences that show no change and do not co-precipitate with Dis3L2 (19) but absent in Dis3L2 target sequences. E-value = 0.016.

**Supplemental Figure 5:**
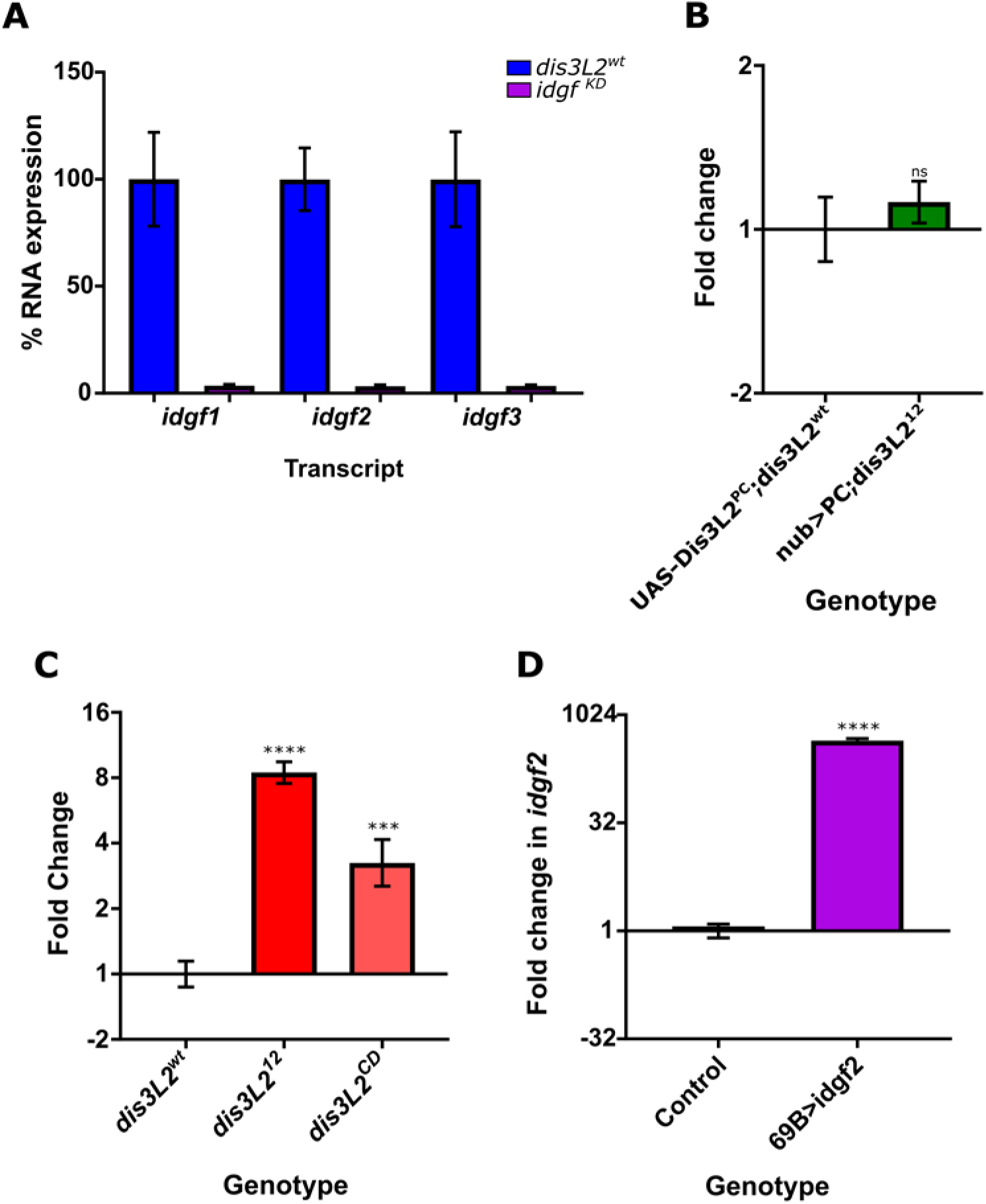
Assessing *idgf2* levels in knockdown, mutant and rescue tissues. **A)** Ubiquitous knockdown of *idgf1*, *idgf2* and *idgf3* by driving specific *UAS-RNAi* constructs with *tub-GAL4* results in >90% knockdown for all targets. n=3, p<0.0004 for all, error bars represent SEM. **B)** Re-expression of Dis3L2 in *dis3L2^12^* wing imaginal discs results in a reduction of *idgf2* mRNA to a level not significantly different from *dis3L2^12^* tissues. n≥6, error bars represent SEM, p=0.4748 (ns). **C)** *idgf2* mRNA is significantly increased in expression in the wing imaginal discs of an independent line carrying a CRISPR generated catalytic dead mutation in the endogenous *dis3L2* locus (*dis3L2^CD^*). n≥4, error bars represent SEM, ****p<0.0001, ***p=0.0006. **D)** Driving UAS-Idgf2 with 69B-GAL4 results in a significant increase in *idgf2* mRNA (*UAS-idgf2/+ ; 69B-GAL4/+*) compared to controls. Controls include both parental lines. n=6, 95% CI, ****=p<0.0001.

**Supplemental Figure 6.**
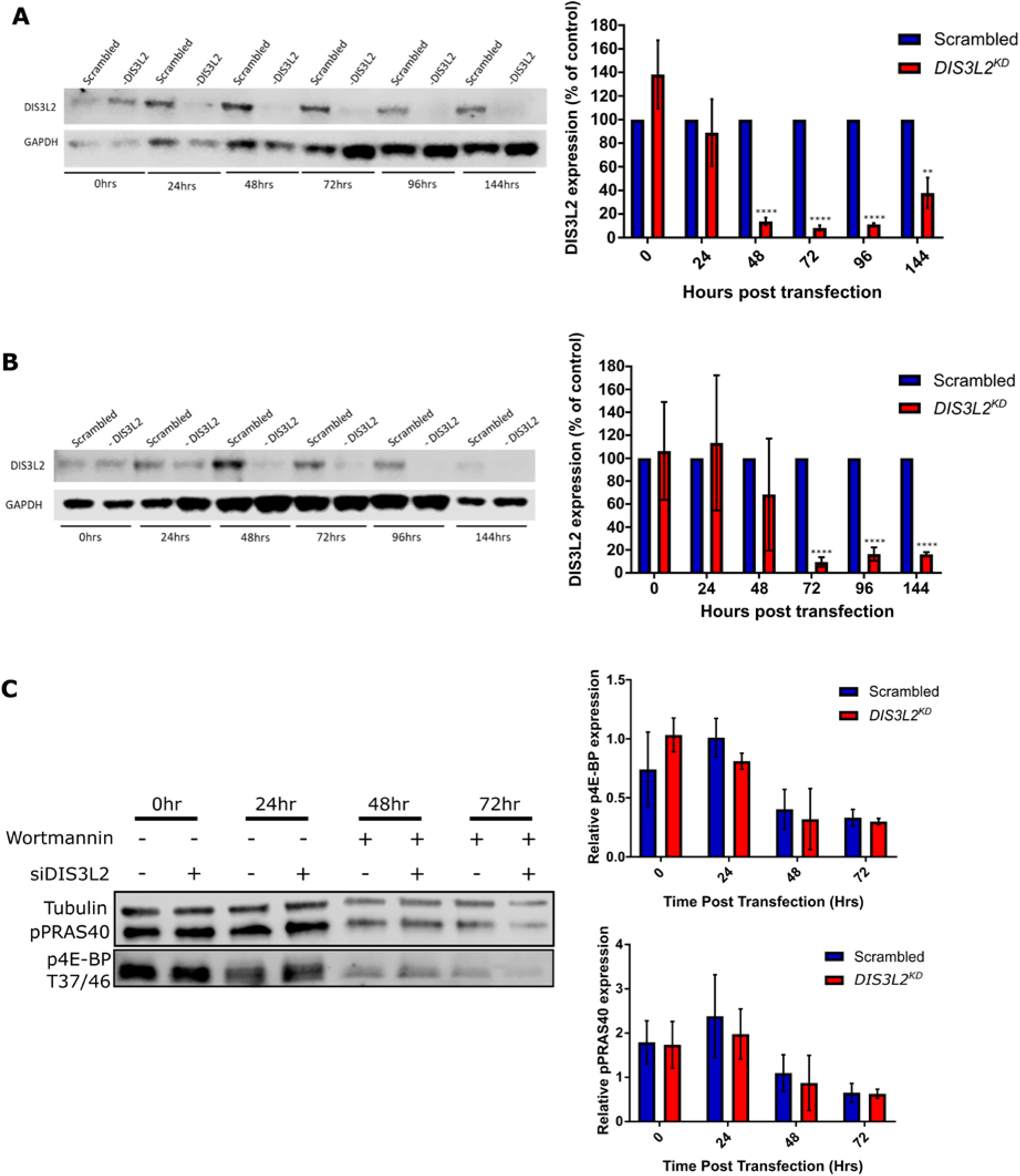
Knockdown of DIS3L2 and validation of Wortmannin activity in human cells. **A)** Knockdown of DIS3L2 is observed 48 hours after transfection and is retained until at least 144hrs post-transfection. Maximal knockdown is observed 72 hours post transfection. n=4, error bars represent SEM, ****=p<0.0001, **=p<0.01. **B)** Knockdown of DIS3L2 is observed 48 hours after transfection and is retained until at least 144hrs post-transfection. Maximal knockdown is observed 72 hours post transfection. n=4, error bars represent SEM, ****=p<0.0001. **C)** Wortmannin treatment reduces pPRAS40 and p4E-BP signal in HEK-293T cells. Quantification of western blots with and without 250nM Wortmannin treatment in DIS3L2^KD^ or Scrambled control HEK-293T cells. n=3, error bars represent SEM.

**Supplemental Figure 7:**
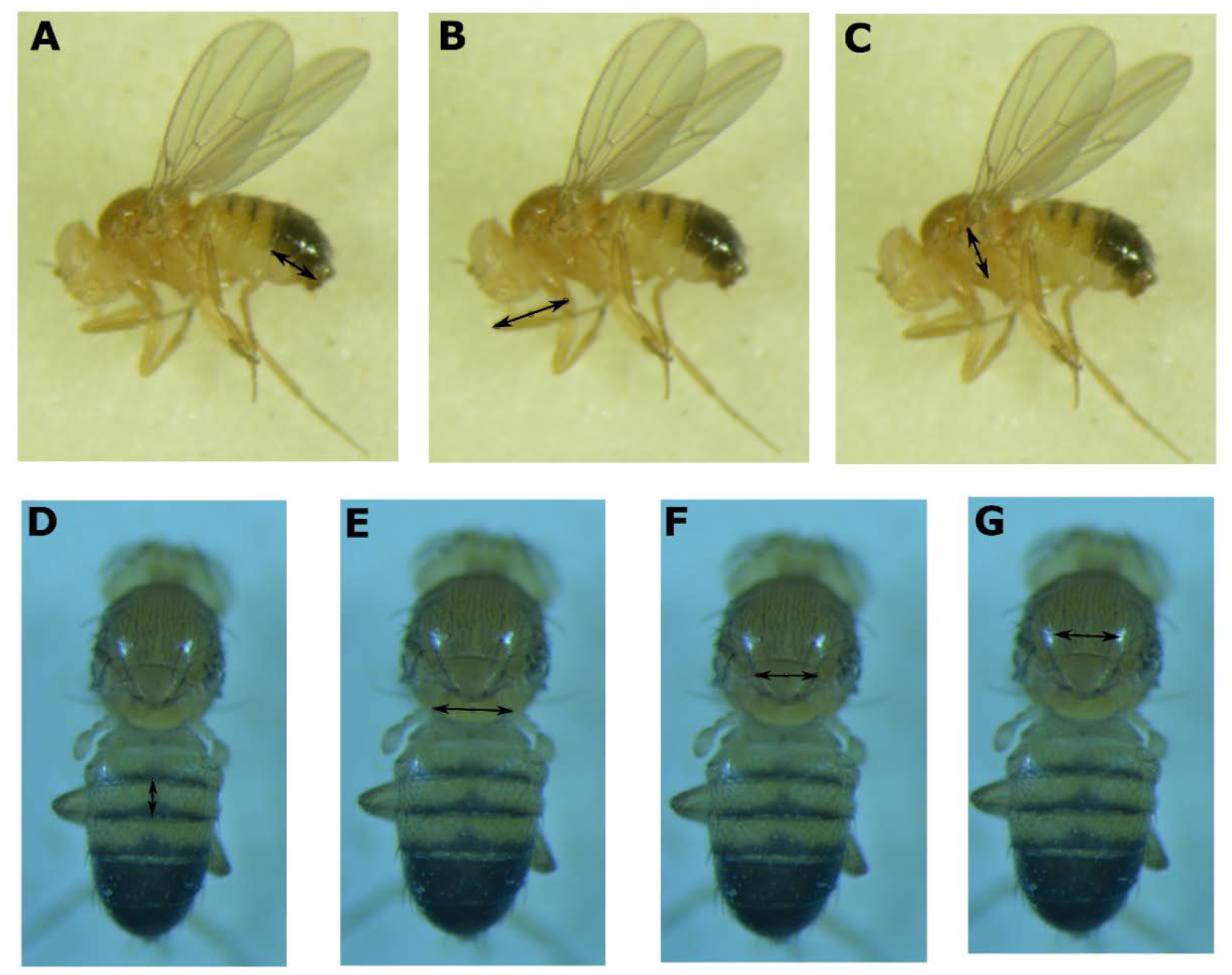
Regions measured to assess male fly size. Measurements taken in ImageJ between the arrows.

**Supplemental Table 1:**
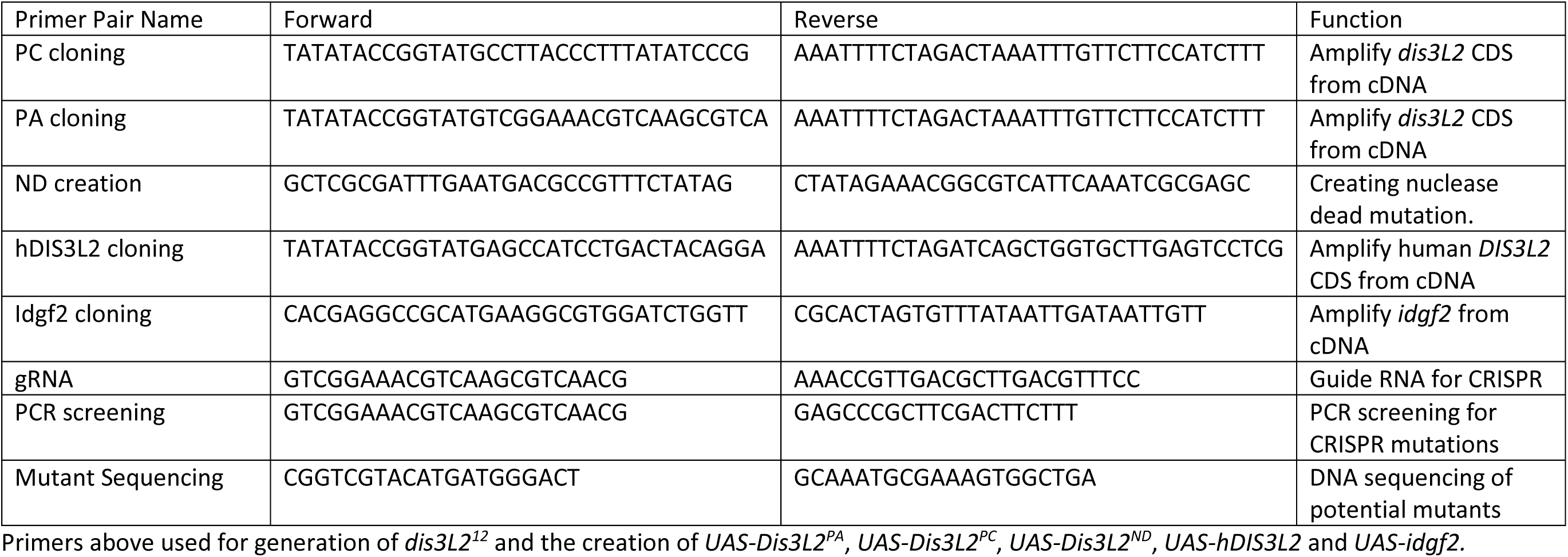
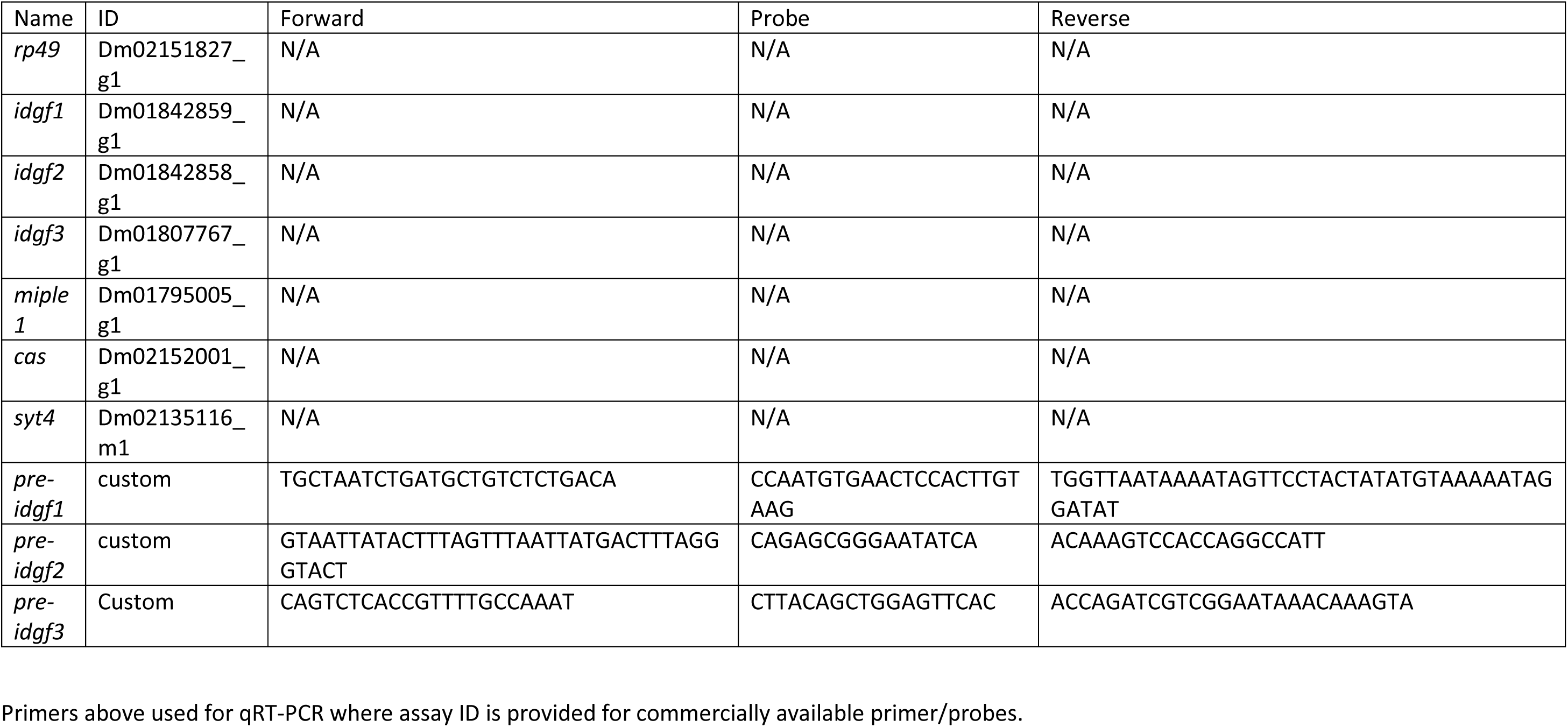
Primers used in this study.

**Supplemental File 1: Additional RNA-seq information**

Summary of read counts and alignments for each RNA-seq replicate. Reads the FlyBase *Drosophila melanogaster* genome (r6.18) using HiSat2 v2.1.0

|  | Replicate | Paired reads | Reads passing filtering | Reads mapped |
| --- | --- | --- | --- | --- |
| *dis3L2<sup>wt</sup>* | 1 | 40640720 | 40287088 | 35223001 |
|  | 2 | 38263277 | 38000207 | 32528177 |
|  | 3 | 37614770 | 37366876 | 31096714 |
| *dis3L2<sup>12</sup>* | 1 | 43538281 | 43265493 | 36918445 |
|  | 2 | 36517562 | 36255393 | 31092625 |
|  | 3 | 39654701 | 39402096 | 33385395 |

Non-default parameters used for RNA-seq alignment and quantification.

| Program | Non-default parameters | Effect |
| --- | --- | --- |
| HiSat2 | -i 20 | Minimum intron size 20bp (default 70bp). |
|  | -l 150000 | Maximum intron size (default 500000). |
|  | -Cufflinks | Adds a tag used by Cufflinks in downstream analysis |
| Cufflinks | -u | More accurate weighting of reads mapping to multiple locations. |
|  | -N | Normalisation to upper quartile of number of fragments mapping to a locus instead of total number of fragments. Improves robustness of differential expression for lowly expressed transcripts. |
|  | -compatible-hits-norm | Only fragments compatible with reference transcript counted in FPKM calculation. |
|  | -M | Supplied a rRNA.gtf file to remove rRNAs from downstream analysis. |
| Cuffquant | -u | More accurate weighting of reads mapping to multiple locations. |
| Cuffdiff | -u | More accurate weighting of reads mapping to multiple locations. |

Comparison between replicates within each genotype

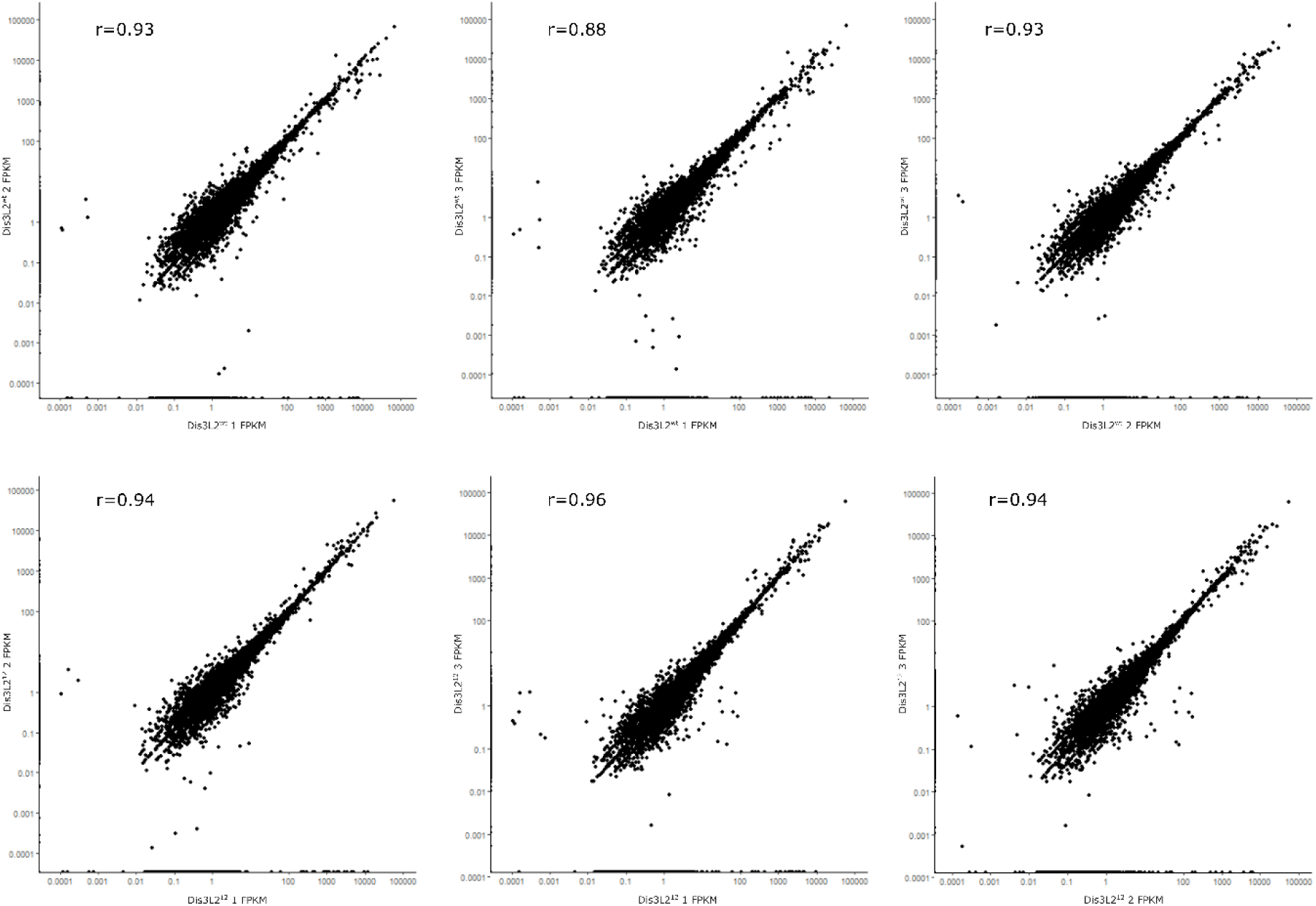

